# Community Curation of Microbial Metabolites Enables Biological Insights of Metabolomics Data

**DOI:** 10.64898/2026.01.24.701521

**Authors:** Helena Mannochio-Russo, Wilhan D. Gonçalves Nunes, Shipei Xing, Fernanda de Oliveira, Andrés Mauricio Caraballo-Rodríguez, Paulo Wender Portal Gomes, Vincent Charron-Lamoureux, Julius Agongo, Nicole E. Avalon, Tammy Bui, Lucia Cancelada, Marc G. Chevrette, Andrés Cumsille, Moysés B. de Araújo, Marilyn De Graeve, Victoria Deleray, Mohamed S. Donia, Mutsawashe B. Dzveta, Yasin El Abiead, Ronald J. Ellis, Donald Franklin, Neha Garg, Harsha Gouda, Claude Y. Hamany Djande, Anastasia Hiskia, Benjamin N. Ho, Chambers C. Hughes, Sunghoon Hwang, Sofia Iliakopoulou, Jennifer E. Iudicello, Alan K. Jarmusch, Triantafyllos Kaloudis, Irina Koester, Robert Konkel, Hector H. F. Koolen, Kine Eide Kvitne, Sabina Leanti La Rosa, Anny Lam, Santosh Lamichhane, Motseoa Lephatsi, Scott Letendre, Sarolt Magyari, Hanna Mazur-Marzec, Daniel McDonald, Ipsita Mohanty, Mónica Monge-Loría, David J. Moore, Thiago André Moura Veiga, Musiwalo S. Mulaudzi, Lerato Nephali, Griffith Nguyen, Martin Orságh, Abubaker Patan, Tomáš Pluskal, Phillip B. Pope, Lívia Soman de Medeiros, Paolo Stincone, Andrej Tekel, Sydney Thomas, Ralph R. Torres, Shirley M. Tsunoda, Fidele Tugizimana, Martijn van Faassen, Felipe Vasquez-Castro, Giovanni A. Vitale, Berenike C. Wagner, Crystal X. Wang, Sevasti-Kiriaki Zervou, Haoqi Nina Zhao, Simone Zuffa, Daniel Petras, Laura-Isobel McCall, Rob Knight, Mingxun Wang, Pieter C. Dorrestein

**Affiliations:** Skaggs School of Pharmacy and Pharmaceutical Sciences, University of California San Diego, La Jolla, CA, USA; Department of Biotechnology, Engineering School of Lorena, University of São Paulo, Lorena, São Paulo State, Brazil; Amazon Integrated Metabolomics Center (CIMAZON), Institute of Natural and Exact Sciences, Federal University of Pará, Rua Augusto Corrêa, 01 - Guamá 66075-110, Belém, PA, Brazil; Department of Pharmaceutical Sciences, School of Pharmacy & Pharmaceutical Sciences, University of California, Irvine, Irvine, CA, USA; Robert A; Center for Marine Biotechnology and Biomedicine, Scripps Institution of Oceanography, University of California, San Diego, La Jolla, CA, USA; Scripps Institution of Oceanography, University of California, San Diego, La Jolla, CA, USA; Department of Chemistry and Biochemistry, University of California, San Diego, La Jolla, CA, USA; Department of Plant Pathology, University of Wisconsin-Madison, Madison, WI 53706, USA; Wisconsin Institute for Discovery, University of Wisconsin-Madison, Madison, WI 53706, USA; Grupo de Pesquisa em Metabolômica e Espectrometria de Massas, Universidade do Estado do Amazonas, 69065-001 Manaus-AM, Brazil; Instituto de Ciências Exatas e Tecnologia, Universidade Federal do Amazonas, 69103-128 Itacoatiara-AM, Brazil; Laboratory of Integrative Metabolomics, Department of Translational Physiology, Infectiology and Public Health, Ghent University, Salisburylaan 133, 9820 Merelbeke, Belgium; Department of Molecular Biology, Princeton University, Princeton, NJ, USA; Research Centre for Plant Metabolomics, Department of Biochemistry, University of Johannesburg, Johannesburg, South Africa; Department of Neurosciences, University of California San Diego, San Diego, CA 92093, USA; HIV Neurobehavioral Research Program, University of California San Diego, San Diego, CA 92093, USA; Department of Psychiatry, University of California San Diego, San Diego, CA 92093, USA; School of Chemistry and Biochemistry, Georgia Institute of Technology, Atlanta, Georgia 30332, United States; Center for Microbial Dynamics and Infection, Georgia Institute of Technology, Atlanta, Georgia 30332, United States; Institute of Nanoscience & Nanotechnology, NCSR Demokritos, Athens, Greece; Department of Microbial Bioactive Compounds, Interfaculty Institute of Microbiology and Infection Medicine (IMIT), University of Tübingen, 72076 Tübingen, Germany; Cluster of Excellence EXC 2124: Controlling Microbes to Fight Infection, University of Tübingen, 72076 Tübingen, Germany; German Center for Infection Research (DZIF), Partner Site Tübingen, 72706 Tübingen, Germany; AquOmixLab, Department of Water Quality Control, Athens Water Supply & Sewerage Company (EYDAP SA), Athens, Greece; National Institute of Environmental Health Sciences, National Institutes of Health, Research Triangle Park, NC, USA; Woods Hole Oceanographic Institution, Woods Hole, MA, USA; Department of Marine Biology and Biotechnology, Faculty of Oceanography and Geography, University of Gdańsk, Gdynia, Poland; Faculty of Chemistry, Biotechnology and Food Science, Norwegian University of Life Sciences, 1432 Ås, Norway; Institute of Biomedicine, Faculty of Medicine, & Turku Clinical Microbiome Bank, Clinical Microbiology & Microbe Centre, Turku University Hospital and University of Turku and Wellbeing Services County of Southwest Finland,Turku, Finland; Department of Medicine, University of California San Diego, La Jolla, CA, USA; Department of Chemistry, Simon Fraser University, 8888 University Drive,Burnaby, Canada; Department of Pediatrics, University of California San Diego, La Jolla, CA, USA; Institute of Environmental, Chemical and Pharmaceutical Sciences, Department of Chemistry, Federal University of São Paulo, Diadema, 09972-270, Brazil; Department of Biochemistry, University of Johannesburg, Johannesburg, South Africa; Institute of Organic Chemistry and Biochemistry of the Czech Academy of Sciences, Prague, Czech Republic; Department of Physical and Macromolecular Chemistry, Faculty of Science, Charles University, Albertov 6, 120 00 Prague 2, Czech Republic; The Centre for Microbiome Research, Queensland University of Technology, 4102, Woolloongabba, Australia; University of Tübingen, Interfaculty Institute of Microbiology and Infection Medicine, Tübingen, Germany; Department of Laboratory Medicine, University of Groningen, University Medical Center Groningen, 9713 GZ Groningen, the Netherlands; Department of Biochemistry, University of California Riverside, Riverside, CA, USA; Department of Chemistry and Biochemistry, San Diego State University, San Diego, California, USA; Center for Microbiome Innovation, University of California San Diego, La Jolla, CA, USA; Department of Computer Science and Engineering, University of California San Diego, La Jolla, CA, USA; Shu Chien-Gene Lay Department of Bioengineering, University of California San Diego, La Jolla, CA, USA; Halıcıoğlu Data Science Institute, University of California San Diego, La Jolla, CA, USA; Hong Kong University of Science and Technology Jockey Club Institute for Advanced Study, Hong Kong SAR, China; Department of Computer Science and Engineering, University of California Riverside, Riverside, CA, USA; Center for Microbiome Innovation, University of California, San Diego, La Jolla, CA, 92093, USA; Collaborative Mass Spectrometry Innovation Center, Skaggs School of Pharmacy and Pharmaceutical Sciences, University of California San Diego, La Jolla, CA, USA; Department of Pharmacology, University of California San Diego, La Jolla, CA, 92093, USA

## Abstract

Microbial metabolites play a critical role in regulating ecosystems, including the human body and its microbiota. However, understanding the physiologically relevant role of these molecules, especially through liquid chromatography tandem mass spectrometry (LC-MS/MS)-based untargeted metabolomics, poses significant challenges and often requires manual parsing of a large amount of literature, databases, and webpages. To address this gap, we established the Collaborative Microbial Metabolite Center knowledgebase (CMMC-KB), a platform that fosters collaborative efforts within the scientific community to curate knowledge about microbial metabolites. The CMMC-KB aims to collect comprehensive information about microbial molecules originating from microbial biosynthesis, drug metabolism, exposure-related molecules, food, host-derived molecules, and, whenever available, their known activities. Molecules from other sources, including host-produced, dietary, and pharmaceutical compounds, are also included. By enabling direct integration of this knowledgebase with downstream analytical tools, including molecular networking, we can deepen insights into microbiota and their metabolites, ultimately advancing our understanding of microbial ecosystems.

## Introduction

Of the thousands of molecules detectable by liquid chromatography-mass spectrometry (LC-MS/MS) in typical biospecimens, the host-associated microbiome modifies 15-70% of them depending on the specific organ or biofluid analyzed^1^. In a typical untargeted metabolomics profile from humans, only about 10% of the acquired spectra can be annotated, and among these, an even smaller portion can be directly traced to microbial origins. Humans have three major sources of microbial metabolites: 1) microbial metabolism of host-derived metabolites^2^; 2) microbial metabolism of molecules from food and beverages^3^; and 3) microbial metabolites assembled *de novo* using proteins encoded by genetic elements often arranged as gene clusters (in bacteria, archaea, fungi, and, recently, discovered to be widespread in phages)^4^. Additionally, microbial metabolites found in humans originate from microbial processing of xenobiotics other than food, such as plasticizers, pollutants, medications, and environmental molecules absorbed through the skin or inhaled ^5–7^.

Despite the critical importance of microbiome-derived metabolites to human health – including those involved in the microbe-gut-brain^8^ and microbe-diet-host axes^9^ – and other ecosystems, there is no centralized knowledgebase where the scientific community can deposit, curate, access, and reuse that knowledge. Existing resources have assessed how the microbiome influences the consumption and production of about 900 largely primary microbial metabolites^10^, and have compiled literature-curated information about 3,269 microbiome-derived metabolites^11^, but most of these metabolites are not unique to microorganisms or have been curated from metabolic models, which tend to capture mainly primary metabolism rather than specialized metabolites which can be more biologically relevant for host-microbiome interactions^12,13^. In addition, some targeted commercial metabolite platforms claim to capture up to ∼140 microbial molecules, but many of those could also come from diet or the host, highlighting the challenge in the field with accurately understanding microbiome-derived metabolites^14^. microbeMASST, our recent tool that enables searching a fragmentation spectra against a reference microbial metabolomics database, allows direct connection between bacteria and fungi and microbially-derived molecules they produce^15^. However, microbial metabolites that have been recently discovered (or even yet to be discovered), the organisms and the genes responsible for their production, and/or their related activities, are not yet systematically cataloged. Therefore, reusing this information is a bottleneck for the community that aims to mechanistically understand the microbiome.

To complement existing microbial metabolite resources, as well as to enable annotation of structurally uncharacterized metabolites (captured as MS/MS spectra), we have created the Collaborative Microbial Metabolite Center knowledgebase (CMMC-KB). Leveraging the Global Natural Product Social Molecular Networking (GNPS)^16^ mass spectrometry analysis ecosystem, the CMMC-KB enables collaboration within the scientific community to curate knowledge on microbial metabolites or metabolites that might influence microbial metabolites (drugs, food, etc). The goal of this initiative is to facilitate biological interpretations of microbiome-derived molecules. With downstream molecular networking integration, the CMMC-KB allows users to visualize MS/MS spectra of compounds classified as microbial metabolites within molecular networks (grouped by MS/MS spectral similarity), even if their structures remain unknown. Furthermore, it provides information on microbial producers, the sources of the molecules, associated genes or sequences, and their biological activities, if known. For a broader investigation of the metabolome, molecules from other sources, such as endogenous molecules, compounds ingested through diet, and drugs, among others, are also included as part of this resource. The CMMC-KB is a user-accessible, collaboratively curated, and continuously evolving microbiome resource. Further, to encourage data deposition, we offer web-based analysis tools, including accessible web applications, that benefit both the data contributors and the broader community. In alignment with the FAIR data principles, we are committed to building this central knowledge hub in collaboration with the scientific community.

## Results and discussion

The CMMC-KB (https://cmmc-kb.gnps2.org/) is a knowledgebase derived from contributions by the scientific community and comprises spectral (MS/MS data) and structural (chemical structures) information about microbially-derived compounds, as well as dietary, host-derived, and other exposure-related compounds. Contributions to the CMMC-KB are facilitated through a dedicated workflow in GNPS2 (a second major implementation of the GNPS ecosystem), enabling users to upload information organized into eight main sections: 1) MS/MS data selection, 2) metabolite identification, 3) taxonomy/phylogeny selection, 4) biosynthesis, 5) activity, 6) references, 7) funding information, and 8) additional comments (**Figure 1a**). The community can contribute to this resource by uploading knowledge for a single molecule at a time or in batches of molecules. A comprehensive documentation page is available to guide users on the recommended inputs (https://cmmc.gnps2.org/deposition_documentation/). While MS/MS spectra are recommended, they are not required, and users may deposit the molecular structure. Additionally, the molecules deposited can be classified as confirmed (e.g., observed experimentally in microbial cultures^17^, observed in colonized but not in germ-free mice, etc.) or predicted to be microbial (e.g., MS/MS of synthetic compounds with matches against other microbial resources like microbeMASST^15^). As of December 2025, the knowledgebase comprises 80,201 MS/MS spectra from 4,998 compounds that were linked to 2,722 microorganisms. These numbers reflect the collective efforts of more than 30 researchers who have contributed to the development of this resource to date^17–21^. Among the compounds deposited, their molecular sources were mainly classified as microbial, drug, or diet-related (**Figure 1b**). The majority of compounds had a known molecular origin, such as drugs, *de novo* biosynthesized by microbes, or diet, with 25.9% classified as unknown/undefined (**Figure 1c**). Since diet, drugs, and host-derived molecules can act as confounders, and because they often influence microbial metabolite production, the CMMC-KB includes and annotates these non-microbial compounds within a single, comprehensive resource.

**Figure 1.**
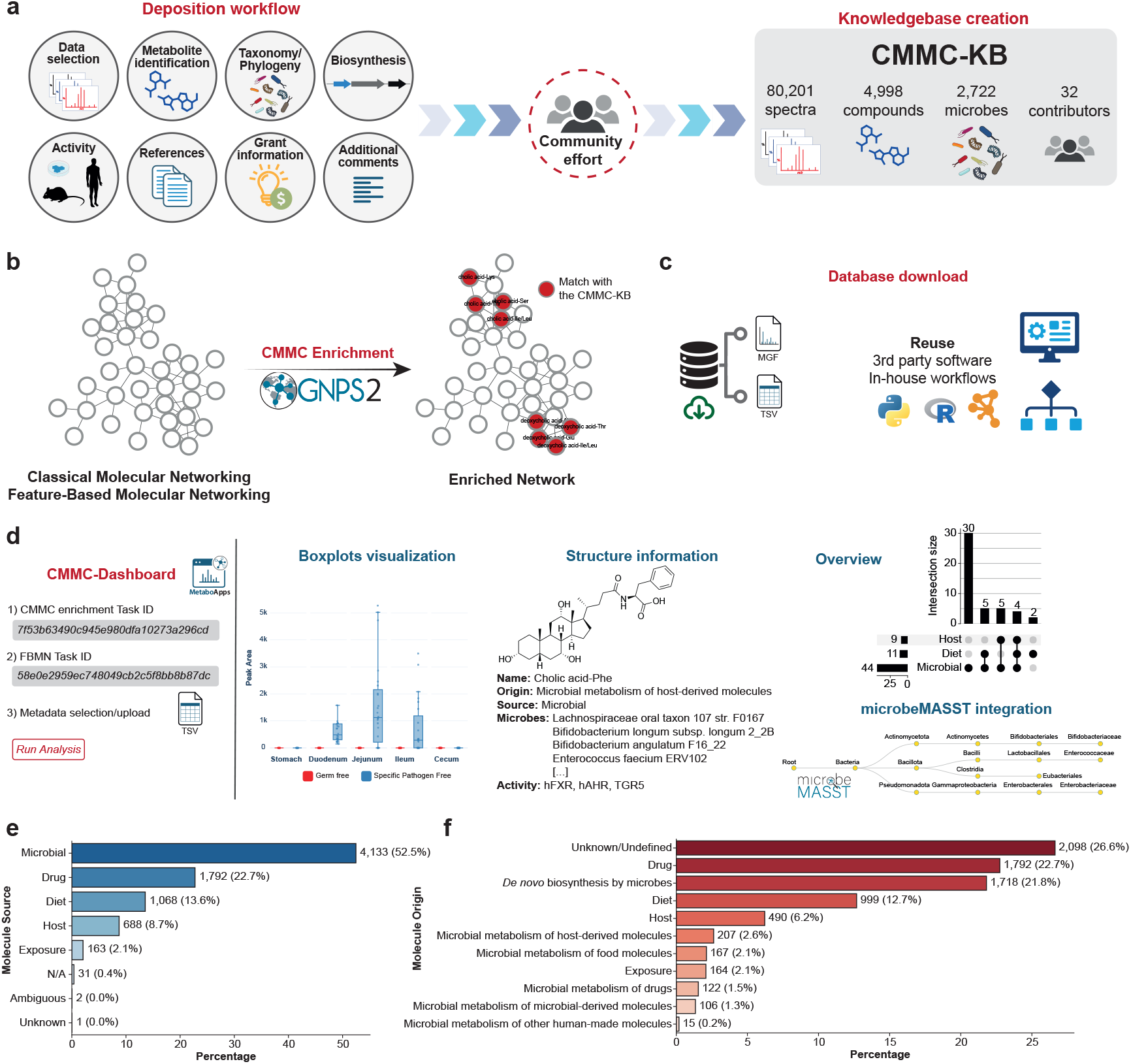
Overview of the CMMC-KB capabilities and depositions. **(a)** Inputs accepted for community depositions and current numbers as of December 2025. **(b)** CMMC enrichment workflow in GNPS2, which annotates molecular networks (generated from Classical or Feature-Based Molecular Networking^25,26^) by matching experimental spectra to the CMMC-KB and retrieving associated metadata. **(c)** Download options as MGF and TSV files, enabling reuse in third-party software and in-house workflows. **(d)** The CMMC-Dashboard is a web application that enables users to utilize outputs from the FBMN and CMMC enrichment workflows, along with uploaded metadata, to generate visualizations for exploring matches to the CMMC-KB (e.g., boxplots for statistical evaluation, structure cards, UpSet-style overviews, and microbeMASST integration). Distribution of the deposited compounds (December 2025) by **(e)** molecule source and **(f)** molecule origin. Icons were obtained from Bioicons.com.

To facilitate the use of information deposited in the CMMC-KB, there are three ways to access and leverage the knowledgebase. First, data are available for direct download in TSV/CSV and MGF formats from the website, allowing integration into customized in-house or third-party workflows. Second, we developed a workflow within the GNPS2 ecosystem that enables downstream enrichment of molecular networks (CMMC enrichment) with information from the CMMC-KB. Finally, we created an interactive web application^22^, CMMC-Dashboard (https://cmmc-dashboard.gnps2.org/), which allows users to visually explore and interpret the data in an accessible and user-friendly manner.

Many compounds deposited as microbial metabolites may also come from other sources. For example, some amino acids and fatty acids can be synthesized by microorganisms, ingested through diet, and also produced by host cells. To address this issue, we refined source annotations in the CMMC-KB by reanalyzing four datasets available in the public domain which contained tissues or biofluids of germ-free (GF) and colonized mice (MSV000079949^1^, MSV000088040^23^, MSV000097485, MSV000090974^24^), also considering mouse diet (chow) for metabolomics data, when available. We ran feature-based molecular networking (FBMN)^25^ followed by CMMC enrichment in GNPS2. Entries initially labelled as “microbial” were selected in the CMMC-Dashboard, and boxplots were plotted for GF vs. colonized mice (and also vs. diet, if available). We defined a metabolite as “microbial-only” when it was absent in the GF group but present in colonized mice (**Supplementary Figure S2a-c**), and added labels as “diet” and/or “host” when the metabolite was detected in GF and/or chow. This classification may include both microbially-produced metabolites and microbe-induced host metabolites, which cannot be distinguished without additional experimental validation. This targeted curation expanded the information available in the CMMC-KB by providing additional classifications for 88 metabolites (1.76% of the compounds deposited).

To illustrate how the CMMC-KB can benefit researchers, we used this resource to investigate microbial metabolites in a subset of the American Gut Project (n = 1,993 files), a citizen-science cohort with participation open to the general population (primarily from the United States, the United Kingdom, and Australia)^27^. In this example, FBMN was performed, followed by CMMC enrichment to annotate features based on spectral matches to the knowledgebase. The source distribution of matched metabolites revealed a diverse chemical landscape, with compounds classified across multiple categories, including microbial, host-derived, and xenobiotic sources (**Figure 2a**). By overlaying this information onto the molecular network, one can rapidly visualize regions enriched in specific source categories (**Figure 2b**). This network-based visualization facilitates hypothesis generation by revealing which networks of structurally related compounds share common sources. Zooming into specific network regions (**Figure 2c-e**) demonstrates the utility of the tool for detailed exploration of individual molecular families, where users can have an integrated view of the source annotations, structural relationships, and associated metadata for compounds of interest. With such an overview, users can target the investigation of specific classes of compounds with important biological functions. For instance, microbially-derived bile acids play crucial roles in immune regulation,^28^ and have been implicated in conditions ranging from inflammatory bowel disease to metabolic disorders and neurocognitive function^29,30^. Similarly, *N*-acyl lipids serve as signaling molecules involved in immune homeostasis, energy metabolism, and gut-brain axis communication^31,32^. The ability to identify and annotate these metabolite families (along with their potential microbial, dietary, or host origins) enables researchers to formulate targeted hypotheses about microbiome-host interactions and prioritize investigations into specific microbial producers, dietary influences, or disease associations. This analysis exemplifies how the CMMC-KB, combined with molecular networking, provides an efficient workflow to survey complex metabolomic datasets and identify features warranting further mechanistic investigation. Importantly, while the biological roles of bile acids and *N-*acyl lipids in gut-microbiome interactions were previously established, the CMMC-KB workflow enabled their rapid annotation and source classification in the American Gut Project cohort – a process that would have required extensive manual literature curation. This cross-cohort validation demonstrates that known metabolite-microbiome relationships can be efficiently detected across diverse population studies using this framework.

**Figure 2.**
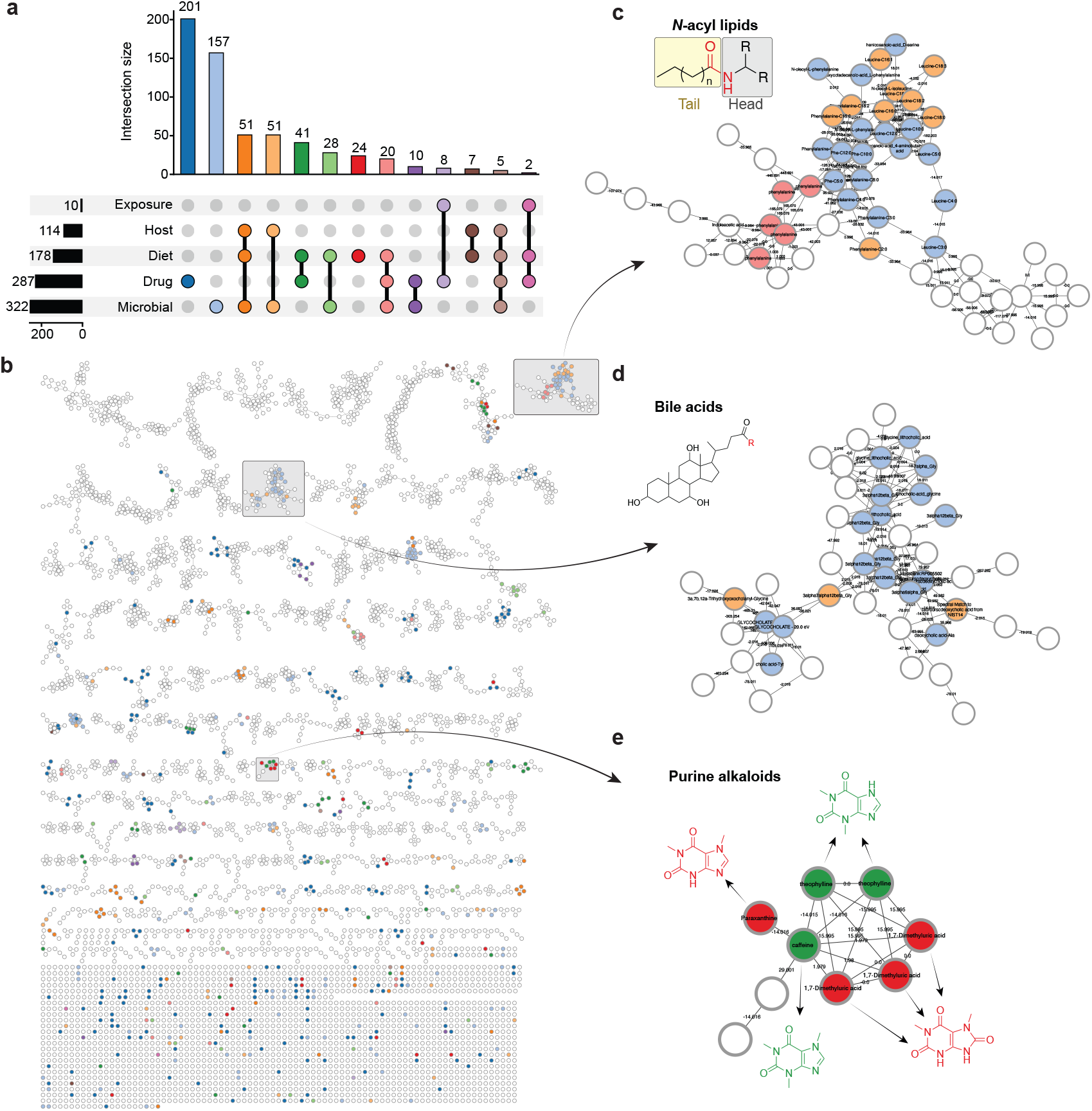
Application of CMMC-KB enrichment to fecal metabolomics data from the American Gut Project. **(a)** Source distribution of metabolites matched to the CMMC-KB from a subset of the American Gut Project (n = 1,993 samples)^27^. The UpSet plot was generated using the CMMC-Dashboard. **(b)** Molecular network visualization with nodes colored by metabolite source annotation from the CMMC-KB. Each node represents a unique mass spectral feature, and edges connect features with similar MS/MS spectra (cosine similarity threshold set to 0.5). **(c-e)** Zoomed-in views of selected molecular networks with distinct source annotations. These subnetworks illustrate the tool’s capability to rapidly identify and visualize structurally related compounds sharing common sources within complex metabolomic datasets. The colors of the nodes in **b-e** match the upset plot colors in **a**.

Beyond this specific use case, the CMMC-KB has been applied to diverse biological contexts that demonstrate its versatility in addressing complex research questions, ranging from the human microbiome, natural products, and environmental fields (**Supplementary Material**). In clinical settings, this resource enabled mapping drug metabolism across multiple biofluids in people with HIV, revealing that while antiretrovirals like ritonavir undergo extensive microbial transformation in the gut, these derivatives remain largely absent from plasma and cerebrospinal fluid (**Supplementary Figure S1**). Comparisons between germ-free and colonized mice facilitated the annotation and the refinement of microbial metabolites, including bile acid conjugates and *N*-acyl lipids, illustrating the dynamic, community-driven nature of the knowledgebase as new data emerge (**Supplementary Figure S2**). Environmental applications include the detection of bioactive cyanobacterial metabolites in Lake Marathon water samples, providing actionable information for water safety management (**Supplementary Figure S3**). In disease contexts, the tool identified a microbiome-derived bile acid conjugate altered by *Leishmania* infection in hamster tissues, linking microbial metabolism to parasite-induced disturbances (**Supplementary Figure S4**). Finally, in coral holobiont research, the CMMC-KB successfully disentangled bacterial versus zooxanthellae metabolic contributions in synthetic communities, revealing siderophore-mediated interactions that would have been difficult to assign using traditional approaches alone (**Supplementary Figure S5**).

When using the CMMC-KB, users should be aware of two key limitations. First, spectral matches are based on cosine similarity^33^ or modified cosine similarity^26^, which cannot easily distinguish isomers that share very similar MS/MS patterns. As a result, isomeric compounds, including those originating from different sources, may have spectra with a high cosine similarity (e.g., deoxycholic acid is a microbial metabolite, chenodeoxycholic acid is host-derived, and their MS/MS cosine similarity is >0.9; **Supplementary Figure S2d**). Consequently, users may obtain spectral matches to metabolites of incorrect biological origin, which highlights the need for follow up experiments and analyses for validation. Whenever possible, users should acquire orthogonal data (e.g. UV-vis, retention time, ion mobility collision cross section (CCS)) obtained from authentic chemical standards for confirmation. Second, the microbial origin of metabolites also requires additional experimental validation beyond spectral matching. Users can employ complementary approaches such as pure culture studies, co-culture experiments with isotope tracing (e.g., ^13^C-labeled substrates), comparisons between germ-free and colonized animal models, or spatial metabolomics to confirm not only the accuracy of the annotation but also its microbial biosynthesis or transformation of the detected compounds. As entries and curated knowledge continue to grow with future studies and depositions, the CMMC-KB will increasingly empower researchers to gain biological insights on the role of the microbiome in human health and diverse ecosystems.

## Acknowledgements

We thank the support from NIH (NIDDK) for the Collaborative Microbial Metabolite Center U24DK133658, support from NSF CAREER award #2047235 to N.G, support from the Research Foundation Flanders (FWO) [V406123N] to M.D.G. The Gordon and Betty Moore Foundation, GBMF12120 and https://doi.org/10.37807/GBMF12120, provided support to P.C.D and A.M.C.-R. This research was supported in part by the National Center for Complementary and Integrative Health of the NIH under award number F32AT011475 to N.E.A.. S.L. was supported by the Research Council of Finland funding (grant no. 363417). T.P. was supported by the Czech Science Foundation (GA CR) grant 21-11563M and by the European Union’s Horizon Europe program (ERC, TerpenCode, 101170268). F.O. was supported by FAPESP (2021/09175-4 and 2022/14603-8). H.H.F.K. was supported by FAPEAM, CNPq (443823/2024-3), and FINEP. L.-I.M. acknowledges the Burroughs Wellcome Fund Investigators in the Pathogenesis of Infectious Disease. R.J.E., D.F.Jr., J.E.I., S.L., and D.J.M were supported by NIH P30 MH062512. R.J.E., D.F.Jr., and S.L. were supported by NIH N01 MH22005 and R01 MH125720. This research was supported in part by the Intramural Research Program of the National Institutes of Health (NIH), National Institute of Environmental Health Sciences (ZIC ES103363). The contributions of the NIH author(s) were made as part of their official duties as NIH federal employees, are in compliance with agency policy requirements, and are considered Works of the United States Government. However, the findings and conclusions presented in this paper are those of the author(s) and do not necessarily reflect the views of the NIH or the U.S. Department of Health and Human Services.

## Methods

### CMMC-KB development

The CMMC knowledge portal was developed using the FAIR (Findable, Accessible, Interoperable, and Reusable) principles as a guideline^34^. It incorporates a series of Python workflows designed to process deposition files and generate visualization tables for all deposited information (**F**indable). In addition, the CMMC-KB server compiles the files required for molecular networking enrichment workflows, including the MGF for the spectral database and structural information for deposited metabolites (**A**ccessible). The KB server provides programmatic access through API endpoints (**I**nteroperable) to download the database files, enabling seamless integration and reuse of information within custom workflows (**R**eusable). The database is automatically compiled daily to ensure the workflows use the most up-to-date information available in the KB.

Each compound with an associated structure in the CMMC-KB is assigned a unique URL, enabling seamless cross-linking to external resources such as NPAtlas.^35,36^ The structure page provides users with tools to explore the molecular structure of metabolites and access all available information for a given molecule and mass spectra available in the knowledgebase. From this interface, users can also contribute additional data by being redirected through a URL to a pre-populated deposition page containing the USI, molecule name, and SMILES/InChI, where further information can be added.

The CMMC-KB Statistics page is a public, daily refreshed summary of the knowledgebase that reports coverage (total unique mass spectra), composition (distributions by metabolite source/origin), and temporal dynamics (new deposits over time), alongside contributor activity.

### CMMC-KB deposition workflow

The CMMC-KB deposition workflow is implemented as a Nextflow-based pipeline^37^ on GNPS2, which runs a series of Python scripts to validate and process user submissions. The deposition workflow supports both single (one molecule) and batch (multiple entries) deposition modes. In single-deposition mode, parameters are provided through a workflow form or YAML file, while in batch depositions, the input is provided through a TSV file. Each entry is checked against controlled vocabularies (e.g., source and origin) and must include valid spectral and structural identifiers: spectra are verified via the Metabolomics Spectrum Resolver API^38^ using USIs, and chemical structures (SMILES or InChI) are validated with the GNPS2 ChemicalStructureWebService API. Following validation, all data is submitted to the CMMC-KB server via POST requests. The necessary templates, including the TSV file and deposition instructions, are fully documented and available at https://cmmc.gnps2.org/deposition_documentation/.

### Network Enrichment workflow

The enrichment workflow is implemented as a Nextflow pipeline available within the GNPS2 ecosystem, and can be launched as a downstream analysis from the Classical or Feature-Based Molecular Networking results. This design enables users to annotate molecular networks with microbial information through a single-click integration. The workflow retrieves molecular networking outputs from both GNPS1 and GNPS2 jobs, including the network (.graphml) and associated spectral (.MGF) files. The retrieved spectra will be matched against the ones available in the CMMC-KB by cosine similarity, and the matches are further enriched with additional metadata if available, including microbial producers, taxonomy, chemical structure, biosynthetic gene clusters, molecular origin, activities, and compound classifications predicted using NPClassifier.^39^ The outputs include a library match TSV table and a new .graphml file with overlaid compound metadata information from the CMMC-KB matches. Additional visualizations, such as the producer lineage and taxonomic distribution, are generated from the NCBI Taxonomy IDs linked to each deposited spectrum. A documentation of the network enrichment workflow is available at https://cmmc.gnps2.org/network_enrichment/.

### CMMC-Dashboard web application

The CMMC Analysis Dashboard (https://cmmc-dashboard.gnps2.org/) was implemented as a web application using the Streamlit Python package to provide interactive access to results from the CMMC-KB enrichment workflow in combination with the FBMN data. The dashboard integrates directly with GNPS2 through Task IDs provided by the user. Task IDs from enrichment and FBMN workflows allow the application to fetch processed files, including enrichment results, FBMN quantification tables, molecular networks, and associated metadata. The dashboard can also be launched directly as a downstream analysis from the enrichment workflow results page, from which the required inputs will be prepopulated in the dashboard interface. A complete documentation for this tool can be found at https://wang-bioinformatics-lab.github.io/GNPS2_Documentation/metaboapp_CMMC_dashboard/.

After the inputs are specified, the dashboard merges enrichment outputs with quantification tables and metadata for downstream analyses. Statistical functionality includes the generation of box plots to compare metabolite abundances across groups, with options for stratification and multiple statistical tests. Overlaps of metabolite sources or origins can be visualized using UpSet plots^40^ derived from the enrichment results. Molecular network exploration is supported through interactive Plotly visualizations that highlight selected features within networks, incorporate delta-mass annotations for network edges, and enable export of figures. The dashboard further integrates microbeMASST^15^, allowing users to perform spectral searches based on a Universal Spectrum Identifier (USI) or feature ID, returning exact or analog matches with compounds from microbial cultures. This allows for taxonomically informed results with corresponding downloadable taxonomic trees.

## Data availability

All the datasets used in this work as use cases of the CMMC-KB are available in MassIVE (massive.ucsd.edu). Raw data files from the American Gut Project, used in Figure 2, are deposited at MSV000080673. The feature finding step was performed in MZmine3, following the previous parameters used for this dataset^19^. Feature-Based Molecular Networking analysis and CMMC enrichment analysis for the American Gut Project use case can be found at https://gnps2.org/status?task=553c08a0e2274572a4edd2ba2d669668 and https://gnps2.org/status?task=2ac40effdb0f404fa6a045a580ff5430, respectively. Additional relevant dataset accessions are provided together with their description in the **Supplementary Material**. Owing to human volunteer protection constraints, the sample metadata for the HIV cohorts will be provided upon request to HNRC: https://hnrp.hivresearch.ucsd.edu/index.php/hnrc-home.

## Code availability

The code used for creating and implementing the CMMC enrichment workflow within the GNPS2 ecosystem is available at https://github.com/Wang-Bioinformatics-Lab/CMMC_GNPSNetwork_Enrichment_Workflow. The code used to create the CMMC-Dashboard MetaboApp is available at: https://github.com/wilhan-nunes/streamlit_CMMC_analysis-dashboard. The code used for dataset analyses can be found at: https://github.com/helenamrusso/CMMC-KB_manuscript.

## Author contributions

H.M.-R., M.W., and P.C.D. conceptualized the project.

W.D.G.N., S.X., and M.W. developed the CMMC enrichment workflow and its integration to the GNPS2 environment.

H.M.-R. and W.D.G.N. developed the CMMC-Dashboard MetaboApp.

H.M.-R., W.D.G.N., V.C.-L., M.D.G. and M.V.F. created documentation.

H.M.-R., W.D.G.N., F.O., A.M.C.-R., P.W.P.G., J.A., N.E.A., T.B., L.C., M.G.C., A.C.M., M.B.A.J., M.D.G., V.D., C.Y.H.D., M.D., M.B.D., K.E.K., Y.E.A., N.G., H.G., B.H., S.H., S.I., T.K., I.K., R.K., H.H.F.K., A.L., S.L., S.L.L.R., P.B.P., M.M.L., S.M., H.M.-M., I.M., M.M.-L., T.A.M.V., M.S.M., L.N., G.N., M.O., A.P., T.P., L.S.M., P.S., A.T., S.T., R.R.T., S.M.T., F.T., M.V.F., F.V.-C., G.A.V., B.W., C.X.W., H.N.Z., S.Z., D.P. contributed to the CMMC knowledgebase.

H.M.-R., W.D.G.N., S.I., R.K., T.K., S.-K.Z., A.H., H.M.-M., L.-I.M., M.M., and N.G. contributed to use cases.

A.K.J., D.M., and R.K. supervised sample handling and acquired data from the American Gut Project cohort.

R.J.E., D.F.Jr., J.E.I., S.L., and D.J.M. developed the clinical cohorts of human immunodeficiency virus (HIV) infection.

W.D.G.N. and M.W. developed the deposition algorithm and the knowledgebase interface.

P.C.D. acquired funding and supervised this project.

H.M.-R., W.D.G.N., and P.C.D. wrote the manuscript. All authors reviewed and edited the manuscript.

## Supplementary Material

## 1. Investigation of microbial metabolism of drugs in human cohorts of people with HIV

### Author: Helena Mannochio-Russo

### Data availability

MSV000092833 (HNRC MIBI, stool samples)

MSV000094927 (CHARTER, cerebrospinal fluid samples)

MSV000096476 (CHARTER, blood plasma samples)

### Feature-Based Molecular Networking jobs

Stool: https://gnps2.org/status?task=0f32c284e015471c96e39bbbefe8f2d2

CSF: https://gnps2.org/status?task=afcb5cdb64c7490ca6862cafd5bbdac1

Plasma: https://gnps2.org/status?task=c249ae6dfbdf4dd384313d5db56c0619

### CMMC network enrichment jobs

Stool: https://gnps2.org/status?task=754604934a5a4730a255abb0c96c7111

CSF: https://gnps2.org/status?task=1594638e5983450aa379584e2bfd274b

Plasma: https://gnps2.org/status?task=d6239466cd8f4207b1b531b0e8aa46c5

Drugs can undergo extensive biotransformations once administered. These transformations may occur through host metabolic processes, such as those mediated by intestine or liver enzymes, or via interactions with the gut microbiome^1^, which collectively encode approximately 150 times more genes than the human genome. Once metabolized, both the parent compounds and their derivatives can circulate throughout the body, potentially impacting multiple tissues and biofluids.

We have previously shown that fecal samples from people with HIV (PWH) harbor a wide range of drugs and drug transformation products^2^. This observation raises important questions about the extent to which these compounds, and especially their microbially mediated metabolites, are distributed across different biofluids. By comparing feces, plasma, and cerebrospinal fluid (CSF) samples from PWH, we aimed to gain a broader understanding of the reach and transformation of commonly used drugs in this population.

To this end, we applied the CMMC-kb enrichment workflow to molecular networks derived from untargeted LC-MS/MS analyses. Fecal samples were obtained from the Microbiome core (MIBI) from the HIV Neurobehavioral Research Center (HNRC), while plasma and CSF samples were sourced from the CNS HIV Anti-Retroviral Therapy Effects Research (CHARTER) study and were paired within individuals. These combined datasets allowed us to survey the molecular diversity of drug-related features across multiple biofluids. We found that drugs commonly used by PWH, such as antiretrovirals, antidepressants, and antipsychotics, were detected across biofluids, but their patterns of transformation and distribution varied considerably (**Supplementary Figure S1**).

Ritonavir, a widely used antiretroviral, was present as a single node in plasma, with no features detected in CSF. In contrast, a large molecular network was observed in fecal samples, including multiple ritonavir analogs. Two of these were annotated and previously shown to be products of microbial metabolism, mediated by “slow grower” and “medium-slow grower” microbial communities. This suggests that microbial transformations of ritonavir are extensive in the gut, but that these derivatives do not enter the bloodstream or central nervous system in detectable amounts. Quetiapine, an antipsychotic used to treat serious mental illnesses such as bipolar disorder, showed a similar pattern. While a broad diversity of analogs appeared in fecal samples, including one with a known microbial transformation signature, only a limited number of features were found in plasma and CSF. Citalopram, a selective serotonin reuptake inhibitor, presented an interesting contrast. In this case, a microbial transformation attributed to a “fast grower” microbial community was observed in feces, but a greater number of analogs were detected in both plasma and CSF. This suggests that while the gut microbiome can modify citalopram, other systemic or host-mediated processes may dominate its metabolism and distribution.

**Supplementary Figure S1.**
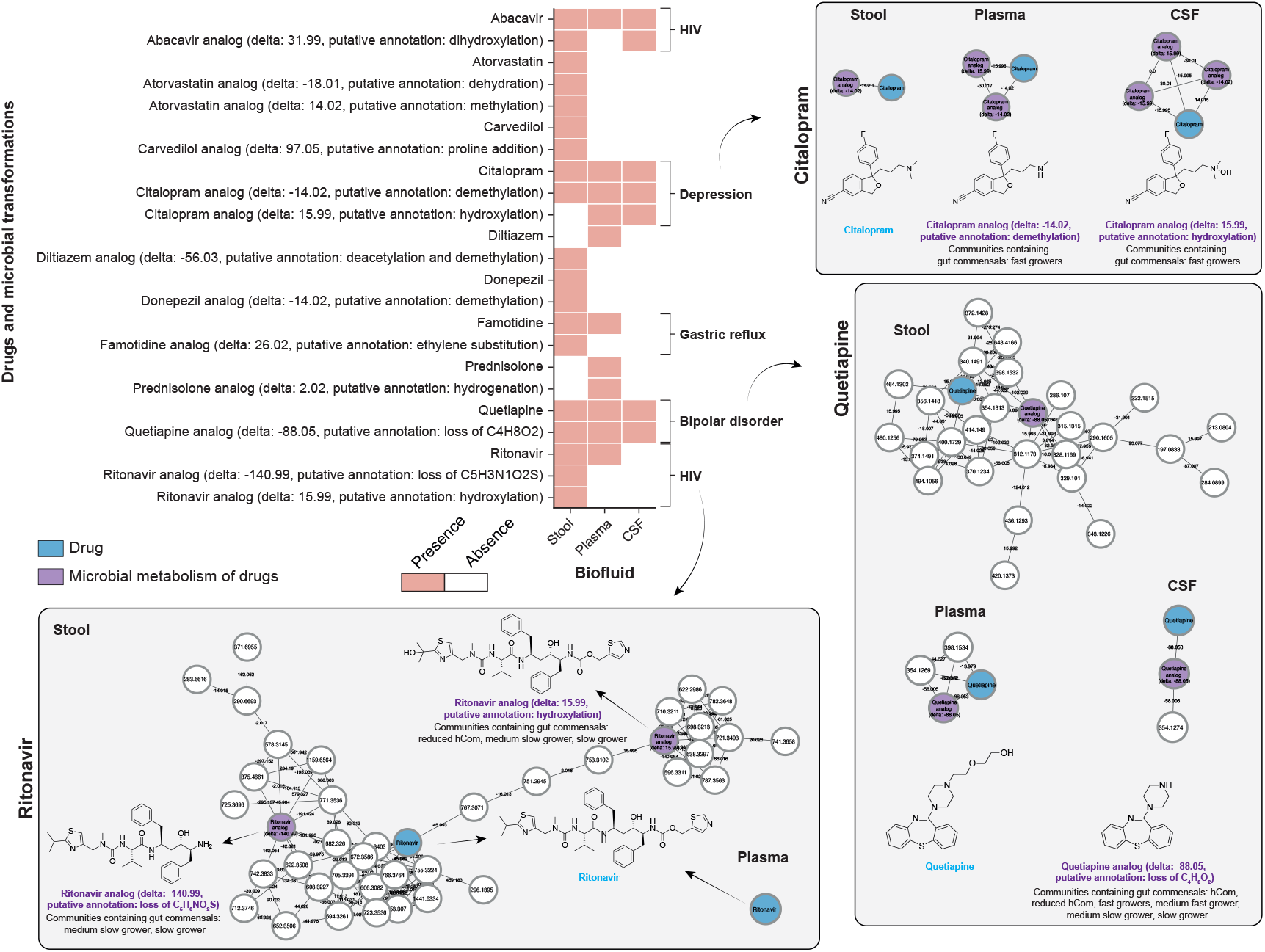
Distribution and microbial transformation of drugs across biofluids in people with HIV. The heatmap shows the presence and absence of drugs and their transformation products across fecal, plasma, and cerebrospinal fluid (CSF) samples from people with HIV. Blue nodes indicate parent drugs, while purple nodes represent products of microbial metabolism. Molecular networks illustrate the structural relationships between parent compounds and their analogs across the three biofluids. Node sizes represent MS/MS spectra, while the edges connect structurally similar compounds based on MS/MS spectral similarity (cosine score > 0.7). The chemical structures that matched the CMMC-KB are representative structures; isomers are possible.

These observations underscore the utility of the CMMC enrichment approach in providing a rapid, scalable framework to visualize and interpret drug metabolism in complex biological matrices. While only a small subset of nodes was annotated with known microbial transformations, many unannotated nodes likely represent previously uncharacterized derivatives. It is important to note that some of the annotated transformations are not uniquely microbial and have also been reported in host-derived metabolism^3^. Further studies are needed to investigate the relative contributions of microbial versus host enzymatic processes in shaping the fate of drugs in the body.

## Methods

The data files from datasets MSV000092833, MSV000094927, and MSV000096476 available in MassIVE were downloaded and further processed in MZmine4 (version 4.1.0, plasma and CSF) or MZmine 3 (version 3.9.0, feces). These datasets were acquired all in the same LC-MS/MS method; however, different extraction protocols were followed. The fecal samples were processed according to swab extractions^4^, while the plasma and CSF samples were extracted using the Phree kit for phospholipid removal. Each dataset was processed individually. The MZmine batch files containing all of the steps and parameters used are available at https://github.com/helenamrusso/CMMC-KB_manuscript. The exported .mgf and .csv files containing spectral and peak area information were used as input in the Feature-Based Molecular Networking Workflow in the GNPS2 environment, and a CMMC enrichment downstream analyses were performed, with the accession links provided above. The parameters used for the molecular networks generation and library searches are available at the provided links.

Each of the CMMC enriched results tables was downloaded and further processed in Python (version 3.7.6) to create the heatmap. Initially, for the stool samples, only HIV+ participants were retained, while for CSF and blood plasma all the participants were HIV+. For each biofluid, only the features classified as “Drugs” or “Microbial metabolism of drugs” as the molecule origin were considered, and the peak area table was transformed to a binary table to represent the presence or absence of a drug or drug metabolite. The heatmap was created using the “seaborn.barplot” package (version 0.12.2), and it was filtered to only contain matches to drugs and drug metabolites of the same drug (i.e., matches to other drugs where no microbial transformation was observed were removed from the figure).

## 2. Body distribution of microbial metabolites in germ-free (GF) versus specific pathogen-free (SPF) mice

### Authors: Helena Mannochio-Russo and Wilhan D. Gonçalves Nunes

### Data availability

MSV000079949

### Feature-Based Molecular Networking job

https://gnps2.org/status?task=94e5dc64e32e4bf381a9030cd198e0d8

### CMMC network enrichment job

https://gnps2.org/status?task=a86fb8dc599d4b66859cf103037efe4a

Germ-free (GF) and colonized specific-pathogen-free (SPF) mice are indispensable models for investigating the impact of the microbiome on host physiology^5^. By providing a controlled system to isolate microbial influences, these models have enabled discoveries that span immune regulation, nutrient metabolism, and xenobiotic transformation^6,7^. Metabolomic profiling of multiple tissues and biofluids from GF and SPF mice has revealed widespread microbial modulation of host biochemistry across all body parts, including the discovery of previously uncharacterized bile acid conjugates^8^. These are powerful models for uncovering both global and tissue-specific chemical signatures of microbial colonization and for exploring the mechanisms through which the microbiome influences health and disease.

In this use case, we re-investigated the dataset in which the bile acids conjugates with phenylalanine, leucine, and tyrosine were initially discovered^8^ to highlight that the use of the CMMC-kb can be a powerful resource to investigate additional microbial metabolites. A Feature-Based Molecular Network job was launched in GNPS2, and a downstream analysis of the CMMC enrichment was done. The CMMC-Dashboard was used to explore the results of this study.

Bile acid conjugates previously reported in the literature for this dataset were successfully annotated and found exclusively in the gastrointestinal tract of SPF mice (**Supplementary Figure S2a**). Additional bile acid conjugates were annotated–arginine, lysine, and serine–and followed the same pattern, further supporting their microbiome-derived source (**Supplementary Figure S2b**). Notably, we also identified *N*-acyl lipids annotated as microbial in origin through the CMMC-kb workflow. These annotations were supported by prior enzymatic studies involving *Faecalibacterium prausnitzii*^*9*^, *in vitro* gut simulations^10^, and by microbeMASST matches using an *N*-acyl lipid library curated from public metabolomics datasets^11^. However, in the present study, *N*-linoleoyl-arginine was also detected in germ-free mice, suggesting that this compound is not exclusively of microbial origin and may also be synthesized by the host^12^. As a result, the annotation in the knowledgebase was updated to reflect both microbial and host sources (**Supplementary Figure S2c**). This case illustrates the dynamic, community-driven nature of the CMMC-kb, which is a living resource that evolves as new data emerges. While entries may initially reflect the state of knowledge at the time of deposition, ongoing research enables continuous refinement and enrichment of the knowledgebase.

The spectral annotations obtained with the CMMC-kb are based on cosine similarity, which cannot reliably distinguish structural isomers since many share MS/MS patterns that result in similarity values above the typical cosine thresholds. For example, deoxycholic acid (microbial origin), and chenodeoxycholic acid (host-derived) yield spectra with close to 0.94 cosine similarity (**Supplementary Figure S2d**). Consequently, annotations should be considered as putative unless supported by orthogonal evidence such as analyses with authentic standards for retention time matching, ion mobility CCS values, etc. _*m/z*_

**Supplementary Figure S2.**
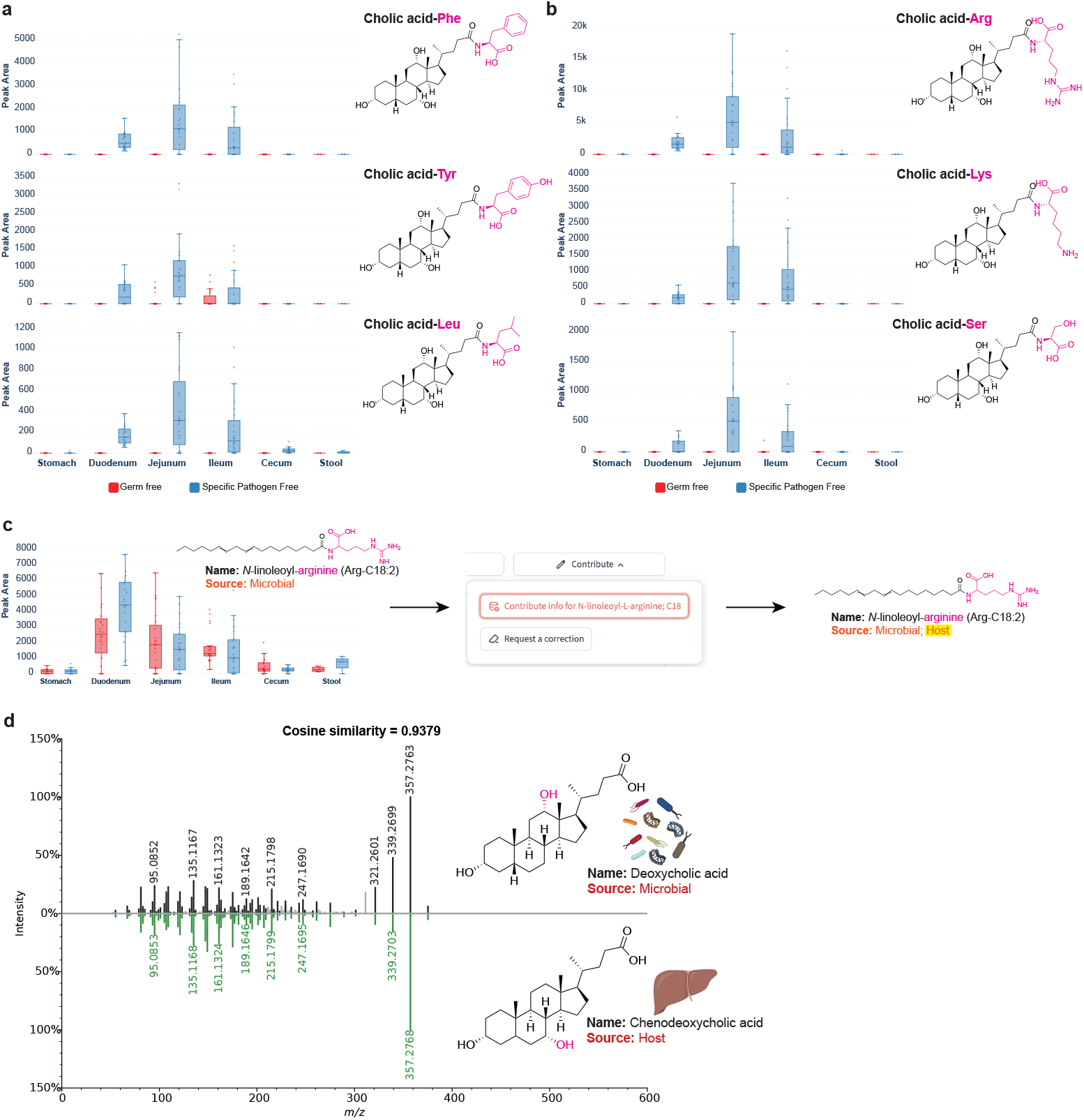
CMMC-kb enrichment reveals microbial and host sources of bile acid conjugates and *N*-acyl lipids in germ-free and colonized mice. **(a-b)** Boxplots showing the tissue distribution of amino acid-conjugated bile acids in the gastrointestinal (GI) tract in germ-free (red) and specific pathogen-free (blue) mice. **(a)** Previously reported bile acid conjugates with phenylalanine, tyrosine, and leucine^8^ are detected in the GI tract of colonized mice. **(b)** Additional bile acid conjugates annotated through the CMMC-kb enrichment, including arginine, lysine, and serine conjugates, follow the same pattern of exclusive presence in SPF mice. **(c)** Arg-C18:2 was initially annotated as microbial based on prior literature reports and microbeMASST matches. Detection in both germ-free and colonized mice prompted a correction in the CMMC-kb, updating the source annotation to include both microbial and host origins. **(d)** Deoxycholic acid (microbial) and chenodeoxycholic acid (host-derived) share highly similar MS/MS fragmentation patterns (cosine similarity = 0.9379), highlighting the need for orthogonal validation methods in cases where isomers have different sources. Icons were obtained from Bioicons.com.

## Methods

The data files from dataset MSV000079949 available in MassIVE were initially downloaded and further processed in MZmine, version 4.0.8. The batch file containing all of the steps and parameters used for this processing is available at https://github.com/helenamrusso/CMMC-KB_manuscript. The exported .mgf and .csv files containing spectral and peak area information were used as input in the Feature-Based Molecular Networking Workflow in the GNPS2 environment, and a CMMC enrichment downstream analysis was performed, with the accession links provided above. The parameters used for the molecular networking generation and library searches are available at the provided links. The CMMC-dashboard was used to inspect the results in the form of boxplots.

## 3. Cyanobacteria metabolites in Lake Marathon

### Authors: Sofia Iliakopoulou, Robert Konkel, Triantafyllos Kaloudis, Sevasti-Kiriaki Zervou, Anastasia Hiskia, and Hanna Mazur-Marzec

### Data availability

MSV000100269

### Feature-Based Molecular Networking job

https://gnps2.org/status?task=306d36d75bf74ae8a0db60db4a15366e

### CMMC network enrichment job

https://gnps2.org/status?task=3b9f9058567b4084b573ae9150d27eee

### CMMC Dashboard link

https://cmmc-dashboard.gnps2.org/?cmmc_task_id=3b9f9058567b4084b573ae9150d27eee&fbmn_task_id=306d36d75bf74ae8a0db60db4a15366e

Cyanobacteria are ancient photosynthetic prokaryotes that are widely distributed in the biosphere^13^. They play major roles as primary producers, in food webs and in biogeochemical carbon, nitrogen, and phosphorus cycles. They can be found in diverse aquatic and terrestrial habitats, such as freshwater bodies, oceans, soil, rocks and microbial mats^14^. Cyanobacteria produce a plethora of over 3,000 secondary metabolites reported so far in the literature, with peptides being the largest compound class^15–17^. Cyano-metabolites have a wide variety of chemical structures and biological effects^18,19^. Several metabolites are toxic to plants, invertebrates, vertebrates, and mammals, including humans^20^. Based on their toxicity, cyanotoxins can be classified into five main groups: hepatotoxins, neurotoxins, cytotoxins, and dermatotoxins^20^. The main routes of exposure to cyanotoxins are drinking water consumption, contact with water during recreational activities, food consumption and inhalation of aerosols near water bodies affected by toxic cyanobacteria^21^. The effects of cyanotoxins can be magnified in food chains via bio-accumulation^22,23^. Cyanobacteria can form blooms resulting in serious water quality problems, such as depletion of oxygen, water taste and odor problems, and the production of cyanotoxins, impacting aquatic and terrestrial wildlife and human health^24^. Cyanobacterial blooms are currently considered a major worldwide threat to public health and aquatic ecosystems^25^. There are indications that the frequency and intensity of cyanobacterial blooms are increasing due to eutrophication, rising carbon dioxide levels, and global warming^26–28^.

Over the last decades, researchers have been focusing on developing and optimizing analytical methods for the detection and structural elucidation of cyano-metabolites in cyanobacteria cultures and environmental samples. Additionally, research has been conducted to assess their bioactivities and to develop sustainable and effective strategies for preventing and controlling toxic cyanobacterial blooms^19,29,30^. Despite the large number of cyanobacteria secondary metabolites reported in literature, certified reference standards are commercially available only for a small number (<20), while MS/MS spectra of cyano-metabolites in public databases are scarce. There is currently no tool to automatically annotate cyano-metabolites and provide information about their properties and producing species, especially in complex environmental samples such as lake blooms, fish, and benthic microbial mats. Microbial interrogation of these complex samples via the workflows integrated into the CMMC knowledgebase (CMMC-kb) would provide a deeper understanding of the microbial dynamics, enabling the identification of compounds that are markers for (toxic) cyanobacteria. In this context, the use of the CMMC-kb could provide insights into ecological and public health management strategies.

The functionality of the CMMC network enrichment workflow was demonstrated by the analysis of a historical sample of Lake Marathon, a landmark artificial water reservoir located in the northeastern part of Attica (Greece), covering an area of 2.45 km^2^. In the early 20th century, Lake Marathon served as the primary water supply reservoir of Athens; currently, it remains connected to one water treatment plant, as a backup water supply. In the past, occurrences of phytoplankton blooms were observed, and cyanotoxins were detected in the lake^31^. In this study, a historical sample collected during a bloom in October 2010 was analyzed by untargeted LC-MS/MS to investigate whether cyanobacterial or other microbial metabolites could be detected. This analysis aimed to enhance our understanding of the microbial composition and the metabolic processes occurring within the lake.

Molecular networking^32^ was employed to explore the chemical space within the sample and to perform library searches based on MS/MS spectra using the GNPS2 environment^33^. The Feature-Based Molecular Networking (FBMN)^34^ workflow resulted in 95 annotations based on MS/MS spectra. Downstream analysis using the CMMC workflow enriched this list, allowing the annotation and prioritization of features specifically associated with cyanobacteria and their neighboring nodes, which were hypothesized to represent possible transformation products or structurally related compounds. **Supplementary Figure S3** highlights the annotated microbial compounds with node coloring representing the producing microbial genera.

**Supplementary Figure S3.**
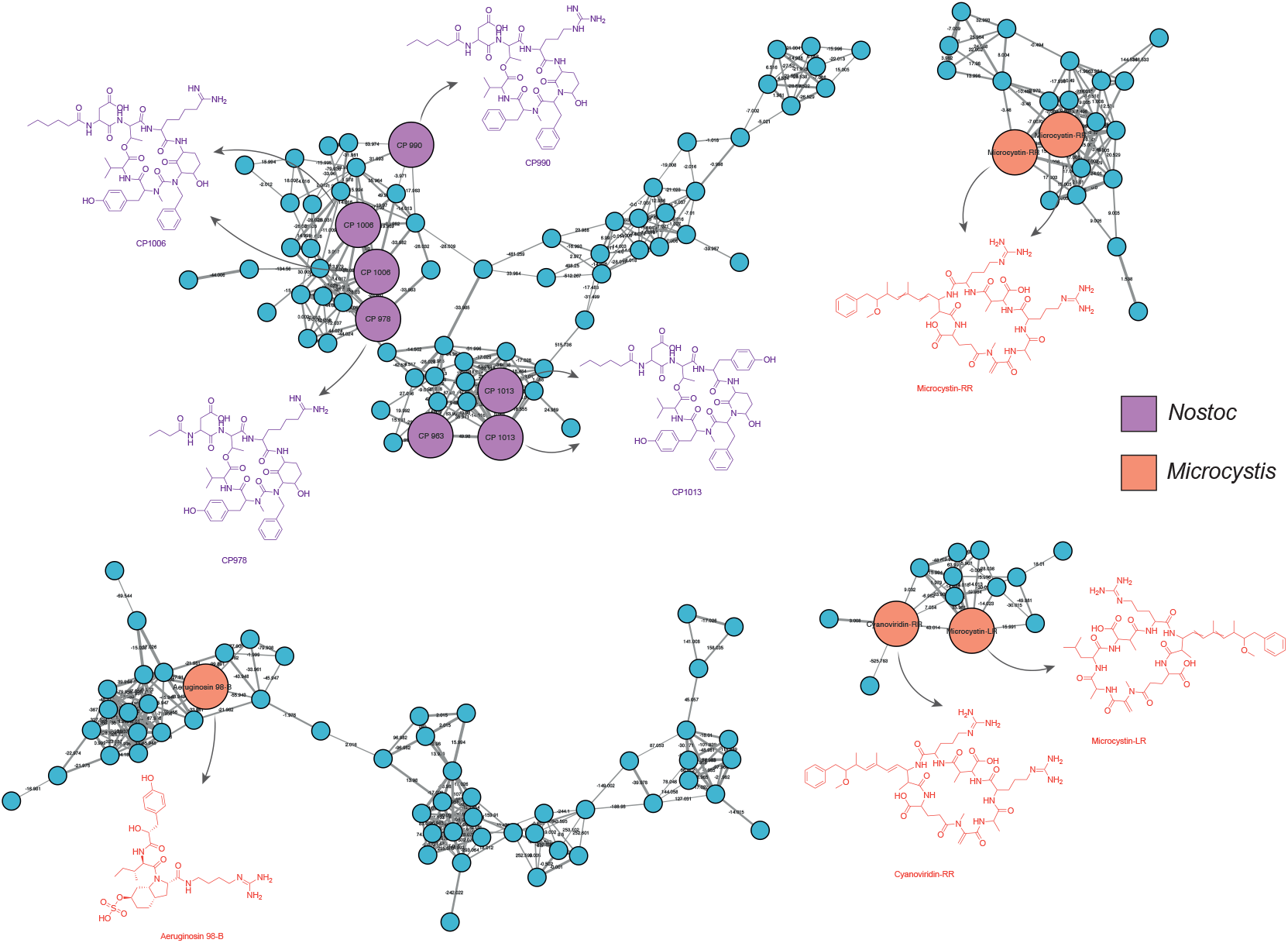
CMMC-kb annotation of cyanobacterial metabolites in Lake Marathon water sample. Molecular network of metabolites detected in a water sample from Lake Marathon, Greece, collected during a cyanobacterial bloom (October 2010). Nodes are colored by producing genus: purple (Nostoc), orange (Microcystis), and blue (unannotated). Four major molecular families were annotated, including cyanopeptolins produced by *Nostoc*, and microcystins and aeruginosins from *Microcystis*. Representative structures are shown for each metabolite class. Each node represents an MS/MS spectrum, while the edges that connect them represent the MS/MS fragmentation similarity (cosine >0.7), and larger nodes represent MS/MS matched with the CMMC-kb.

Features linked to cyanobacteria species were annotated, raising concerns due to their known roles as primary producers of bioactive metabolites, such as microcystins, cyclic heptapeptides that exhibit significant hepatotoxicity to humans and animals^20^. Specifically, two of the most prevalent microcystins (MC), MC-LR and MC-RR, were annotated by the CMMC-kb. These compounds were grouped in a network alongside ten nodes, which were hypothesized to belong to the same molecular family of MCs. Furthermore, several cyanopeptolins (CP), a class of bioactive depsipeptides recently characterized from *Nostoc edaphicum* CCNP1411^35^, were also annotated. Cyanopeptolins are produced by cyanobacteria such as *Planktothrix, Microcystis, Anabaena* sp. and include over 300 congeners with inhibitory activity against trypsin and chymotrypsin, as well as potent cytotoxic effects on HeLa cells^15,29,35^. The CP group represented one of the largest networks in the analysis, with five annotated congeners (CP1013, CP963, CP990, CP1006, CP978). Aeruginosin 98-B was annotated in a network with 56 nodes. Aeruginosins are nonribosomal linear tetrapeptides with inhibitory activity against serine proteases, produced by cyanobacteria such as *Microcystis, Nodularia spumigena, Oscillatoria/Planktothrix, Nostoc*, and *Aphanizomenon*^*36,37*^.

The findings provided by the CMMC-kb enrichment of the molecular network are important for the assessment of the chemical and biological status of the water body. They reveal the occurrence of cyanobacteria, and among them, cyanotoxin producers. In this case, because Lake Marathon serves as an alternative water source for human consumption, the findings support actions by the water supplies to assess the related risks in the context of Water Safety Plans (WSP).

## Methods

### 1.1 Sample Preparation

The original water sample (1 L) was collected in October 2010 from Lake Marathon (Greece), immediately processed (filtered through a glass-fiber filter, freeze-dried), and stored at room temperature. The sample was extracted twice with 1 mL methanol : water (3:1 v/v) with a vortex (Auxilab 681/22, Spain). The extracts were sonicated for 15 min in an ultrasonic water bath (DU-32 DUC, Argo Lab, Germany). The combined extract was centrifuged for 10 min at 10000 rpm and ambient temperature (Heraeus Biofuge 13, Germany). The supernatant was evaporated to dryness under a nitrogen stream (N-EVAP, Organomation, USA). The residue was re-dissolved in 250 µL of a methanol-water mixture (1:4 v/v) and sonicated for 5 min. The resulting solution was analyzed by LC-HRMS

### 1.2 LC-HRMS data acquisition

Analysis was carried out using liquid chromatography coupled with quadrupole-time of flight mass spectrometry (LC-qTOF). An Elute HPG1300 HPLC system (Bruker Daltonics) equipped with an Atlantis T3 C18 column (100 Å, 3 µm, 2.1 mm X 100 mm, Waters) at 30 °C were used. Mobile phases were water (A) and acetonitrile (B) both acidified with 0.1% formic acid. The HPLC gradient elution was: 25% B (held for 2 min), 25-100% B within 18 (held for 3 min) and 100-25% B within 1 min. Re-equilibration time was 6 min after each run. The flow rate was 0.2 mL/min and the injection volume was 10 μL. An Impact II qTOF (Bruker Daltonics) equipped with an electrospray ionization source (ESI) in positive mode was used, operating in AutoMS (DDA) mode. The ESI and MS parameters were: capillary voltage 3100 V, nebulizer gas 1.0 bar, dry gas 6.0 L × min^−1^, dry gas temperature 220 ℃, hexapole 100 Vpp and pre-pulse storage 8 μs. Stepping mode was activated as follows: collision RF from 250 Vpp to 700 Vpp (50–50% of the timing), transfer time from 20 μs to 80 μs (50–50% of the timing) and collision energy from 40 eV to 60 eV (50–50% of the timing). Full scan accurate mass spectra were obtained in the range 50– 1300 *m/z* in AutoMS with dynamic exclusion. Mass calibration was carried out in every sample run using sodium formate cluster ions (10 mM). Bruker’s HyStar and Data Analysis software was used for data acquisition, calibration, and raw data conversion to .mzXML format before further processing. The sample, a procedural blank and a reference standard solution containing 14 cyanobacterial peptides were analyzed in duplicate.

### 1.3 Data processing, molecular networking and CMMC workflows

Mass spectrometry data were pre-processed in MZmine 4.1.0 for MS feature detection, deconvolution, alignment, deisotoping and gap filling. The exported files (quantification table in .csv format and MS^2^ spectra file in .mgf format) from MZmine and a sample metadata file were exported for the FBMN workflow.

Molecular networks were built using the FBMN workflow in GNPS2. The imported data were filtered by removing all MS/MS fragment ions within ±17 Da of the precursor *m/z*. MS/MS spectra were window filtered by choosing only the top 6 fragment ions in the ±50 Da window throughout the spectrum. The precursor ion mass and MS/MS fragment ion tolerances were set to 0.02 Da. Network edges were filtered to have a cosine score above 0.6 and more than 4 *m/z* matched peaks. Edges between two nodes were kept in the network only if each of the nodes appeared in each other’s respective top 10 most similar nodes. The maximum size of a molecular family was set to 100, and the lowest scoring edges were removed from molecular families until the molecular family size was below this threshold. Matching of spectra included in the molecular networks against GNPS2 spectral libraries was carried out. Matches were filtered for cosine scores above 0.7 and at least 5 matched peaks. Molecular networks were visualized in Cytoscape.

The CMMC enrichment downstream analysis of FBMN was carried out in GNPS2 using the same parameters as in the FBMN workflow.

## 4. Amino acid-conjugated bile acid impacted by *Leishmania* parasite infection

### Author: Laura-Isobel McCall

### Data availability

MSV000085991

### Feature-Based Molecular Networking jobs

https://gnps2.org/status?task=bd111efa23e5441686e570f388ed8910

### CMMC network enrichment jobs

https://gnps2.org/status?task=0d818f0dbe4b46fa982bb14ebdfe61aa

### CMMC Dashboard link

https://cmmc-dashboard.gnps2.org/?cmmc_task_id=0d818f0dbe4b46fa982bb14ebdfe61aa&fbmn_task_id=bd111efa23e5441686e570f388ed8910

*Leishmania donovani* parasites are the causative agents of visceral leishmaniasis, a disease that manifests with hepatomegaly, pancytopenia, and fever^38^. There is increasing evidence that the microbiota plays a causal role in visceral leishmaniasis pathogenesis^39^. To identify metabolites of microbiota origin and impacted by *L. donovani* infection, we re-analyzed the feature-based molecular networking data from a previous metabolomics analysis of the liver, gut, and spleen from *L. donovani-*infected and antibiotic-treated or untreated hamsters^39,40^, using the CMMC enrichment workflow and filtering for metabolite features annotated as being of exclusively microbial origin. Noticeably, one of these annotations was alanine-conjugated cholic acid. Bile acids are regulators of serum lipid levels^41^. Analysis of the liver metabolome in *L. donovani-* infected hamsters previously revealed changes in multiple primary and secondary bile acids^40^. Using the CMMC-kb dashboard, we visualized the peak areas of alanine-conjugated cholic acid across experimental groups in the liver and observed that this metabolite was increased in infected livers, and that its levels were abrogated upon antibiotic treatment (**Supplementary Figure S4**). These results can help explain mechanisms of metabolic alteration in visceral leishmaniasis, particularly the observed disturbances in lipid metabolism^42^.

**Supplementary Figure S4.**
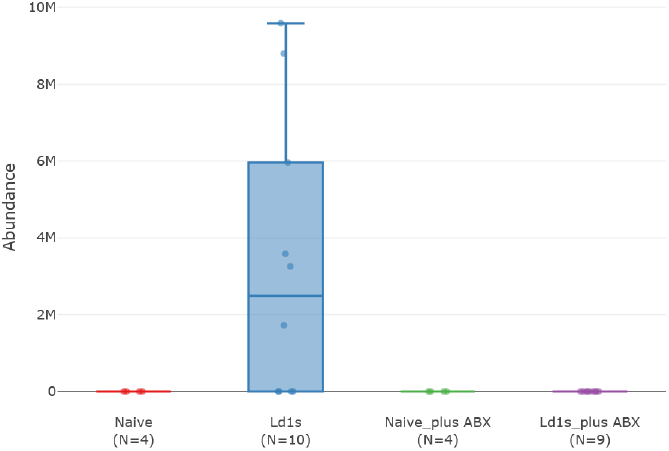
CMMC-kb dashboard visualization of alanine-conjugated cholic acid across experimental groups in the liver. Ld1S, *L. donovani* strain 1S-infected animals. ABX, antibiotics.

## Methods

Sample preparation, metabolite extraction, LC-MS data acquisition, and MZmine data processing were described in Lesani *et al*^*40*^. Feature-based molecular networking and CMMC enrichment analysis workflow were performed using GNPS2, and the parameters used can be accessed in the job links.

## 5. Metabolomic profiling of coral symbiont synthetic communities

### Authors: Mónica Monge-Loría, Neha Garg

### Data availability

MSV000098895

### Feature-Based Molecular Networking jobs

https://gnps2.org/status?task=f1a34201d028428e89aea0e2388f08bf

CMMC network enrichment jobs

https://gnps2.org/status?task=b98e11315ef14aec9e7260b2da7e43c6

### CMMC Dashboard link

https://cmmc-dashboard.gnps2.org/?cmmc_task_id=b98e11315ef14aec9e7260b2da7e43c6&fbmn_task_id=f1a34201d028428e89aea0e2388f08bf

Corals host a variety of prokaryotic and eukaryotic organisms, collectively known as the coral holobiont^43^. The metaorganism incorporates dynamic populations of bacteria, archaea, fungi, viruses, and zooxanthellae. The tripartite relationship between corals, zooxanthellae, and bacteria has been proposed to shape the holobiont’s health and stress tolerance^44,45^. However, while host-zooxanthellae and host-bacteria interactions have been extensively studied^46–50^, the relationship between these two symbionts remains largely underexplored. Their interactions are proposed to be modulators of zooxanthellae productivity and bacterial community composition, ultimately impacting the collective holobiont^51^.

To better understand how these holobiont members interact, we have cocultured coral-isolated zooxanthellae and bacteria. Single-vessel cocultures pose challenges for monitoring population densities and community composition^52^, particularly when culturing organisms with disparate growth rates, such as zooxanthellae and bacteria. A membrane-separated coculture approach overcomes this barrier, additionally preventing antagonistic effects from cell-to-cell contact. This setup prioritizes interactions mediated by chemical signaling, amenable to untargeted metabolomics profiling. Although spatial partitioning improves the stability of the coculture^53^, identifying the precedence of the detected metabolites in liquid media remains a challenge^54,55^. The ability to identify microbially-derived metabolites with the CMMC enrichment analysis allows us to disentangle the metabolomic profile of complex synthetic communities.

In this study, we performed the CMMC enrichment downstream analysis on an FBMN generated for cocultures between *Breviolum* sp. zooxanthellae and three coral-associated bacteria. This analysis identified 38 features from microbial sources, differentially produced under the diverse culture conditions (**Supplementary Figure S5a**). For features such as andrimid and moiramides, detected in *Vibrio coralliilyticus* Cn52-H1 monocultures and in their coculture with *Breviolum* sp., the CMMC enrichment serves as confirmation of their microbial origin. This analysis becomes increasingly useful for features such as desferrioxamines E and G, uniquely detected in the coculture of *Halomonas* sp. Cn5-12 and *Breviolum* sp. (**Supplementary Figure S5b**). Lack of detection in either monoculture precludes a straightforward biosynthetic attribution. Interestingly, desferrioxamine E was detected in both the upper and lower wells, inoculated with *Halomonas* sp. Cn5-12 and *Breviolum* sp., respectively, whereas desferrioxamine G was exclusively detected in the lower, zooxanthellae-inoculated wells. Given the localization of this metabolite, a traditional analysis could have led us to ascribe the production of desferrioxamines to *Breviolum* sp.. However, the CMMC enrichment match and identification as a microbial metabolite allows us to putatively assign these compounds to *Halomonas* sp. Cn5-12. The exclusive identification of desferrioxamines in coculture evidences the inter-domain interactions between *Breviolum* sp. and *Halomonas* sp. Cn5-12. These compounds are classified as siderophores, small molecules with a high affinity for iron, used by diverse organisms to chelate and incorporate the trace metal^56^. Therefore, their upregulation in coculture highlights the role of iron and siderophores as mediators of ecological relationships within the coral holobiont. The ability to identify metabolites of bacterial origin in synthetic communities helps unravel the biological implications of these chemical interactions.

**Supplementary Figure S5.**
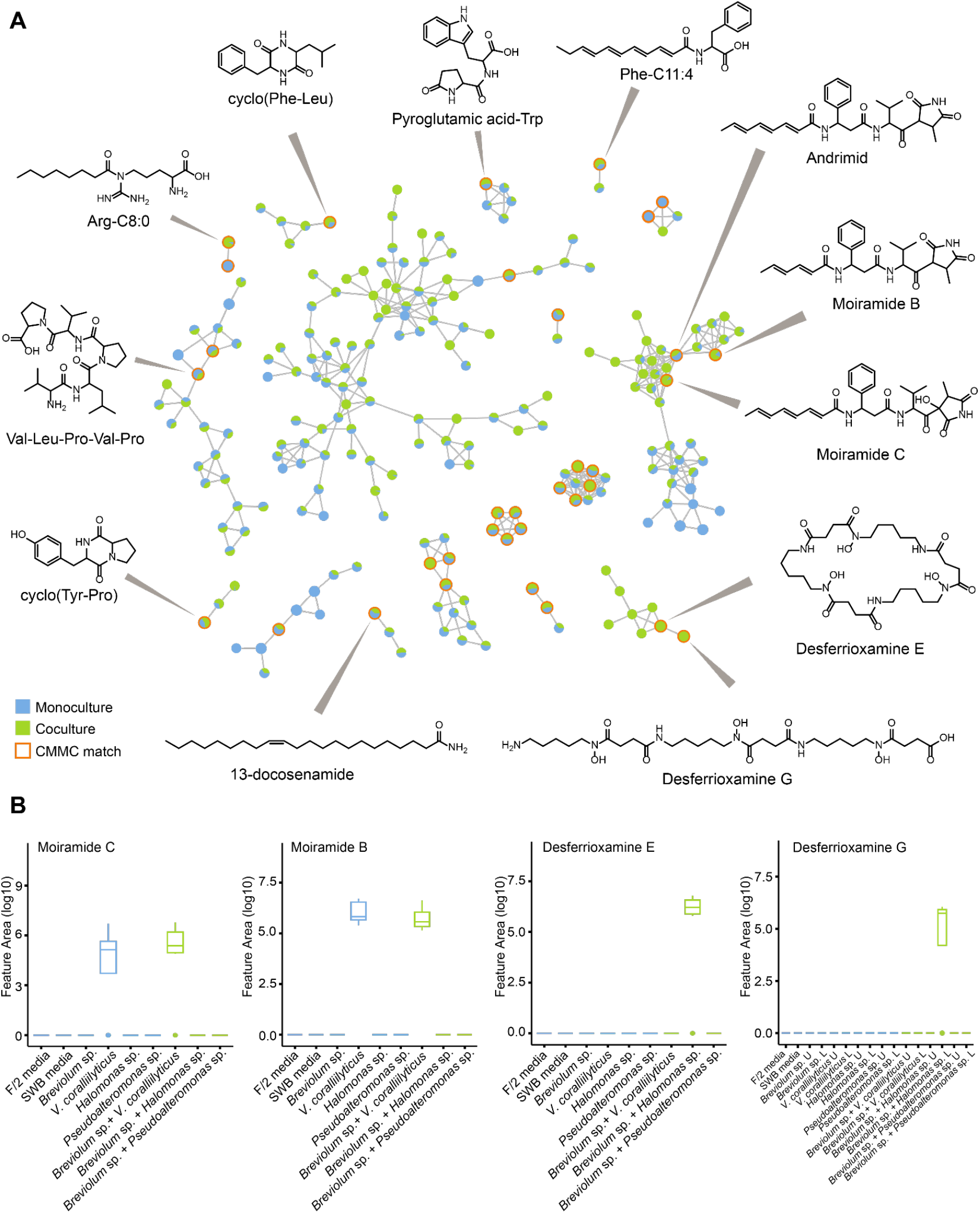
**A)** Representative FBMN for zooxanthellae and bacteria mono- and cocultures. **B)** Boxplots of the relative abundances of moiramides and desferrioxamines in the different culturing conditions. U= upper well, bacterial culture; L= lower well, zooxanthellae culture.

## Methods

### Culturing and sample preparation

Bacterial isolates (*Vibrio coralliilyticus* Cn52-H1, *Halomonas* sp. Cn5-12, and *Pseudoalteromonas* sp. McH1-7) were received from Dr. Valerie Paul. Bacteria were cultured in salt water broth (SWB) and incubated at 30 °C for 24 h while shaking (200 rpm). *Breviolum* sp. zooxanthellae (strain Mf 1.5b, Cp type 184) were received from Dr. Andrew Baker, originally deposited in the BURR collection by Dr. Marie-Alice Coffroth. Zooxanthellae were cultured in Guillard’s media (F/2; Sigma, Burlington, MA) at 21 °C on a 12:12 light cycle. Zooxanthellae were grown for a week prior to inoculation of cocultures. Synthetic communities, as well as the corresponding controls, were cultured on crosstalker plates (Fluidome, Calgary, Canada) for 24 h under the same conditions previously described for zooxanthellae. *Breviolum* sp. was inoculated on a lower, 16-well plate, whereas bacteria were inoculated in an upper, 96-well plate. The crosstalker plates incorporate a semi-permeable membrane that allows for metabolite exchange and chemical interactions. Cultures were harvested and extracted via solid-phase extraction (SPE) using a 25 mg C18 column (Thermo Scientific, Waltham, MA). Analytes were eluted with 20%, 50% and 100% acetonitrile (MeCN), dried *in vacuo* and stored at -20 °C until data acquisition.

### LCMS/MS data acquisition

The dried extracts were resuspended in 200 µL of 100% MeOH containing 1 µM of sulfadimethoxine as an internal standard. The resuspended extracts were analyzed using an Agilent 1290 Infinity II Ultra High-Pressure Liquid Chromatography (UHPLC) system (Agilent Technologies; Santa Clara, CA, USA) with a Kinetex 1.7 μm C18 reversed phase UHPLC column (50 × 2.1 mm) (Phenomenex; Torrance, CA, USA) coupled to an ImpactII ultrahigh resolution Qq-ToF mass spectrometer (Bruker Daltonics, GmbH, Bremen, Germany) equipped with an electrospray ionization (ESI) source. Chromatographic separation was performed with the following mobile phase gradient: 5% solvent B (MeCN, 0.1% (v/v) formic acid) and 95% solvent A (H_2_O, 0.1% (v/v) formic acid) for 3 min, a linear gradient of 5% B–95% B over 17 min, held at 95% B for 3 min, 95% B–5% B in 1 min, and held at 5% B for 1 min, 5% B-95% B in 1 min, held at 95% B for 2 min, 95% B−5% B in 1 min, and held at 5% B for 2.5 min, at a flow rate of 0.5 mL/min throughout. MS spectra were acquired in positive ionization mode from *m/z* 50 to 2000 Da. For MS^2^ data acquisition, the eight most intense ions per MS^1^ were selected for fragmentation. A basic stepping function was used to fragment ions at 50% and 125% of the CID calculated for each *m/z* with timing of 50% for each step. The MS/MS active exclusion parameter was set to two, and the active exclusion was released after 30 s. The mass of the internal lock-mass calibrant was excluded from the MS^2^ list. UV data was acquired with a UV DAD detector (Agilent Technologies) from 190 nm to 400 nm, with a 2 nm step. Zero offset was set at 5% along a 1000 mAU attenuation. Data was acquired throughout the LC run with a >0.1 min peak width.

### Untargeted metabolomics data analysis

Raw data were converted to mzXML format, using vendor proprietary software. Metabolite features were extracted using MZmine 2.53^57^, performing mass detection, chromatogram building, chromatogram deconvolution, isotopic peak grouping, retention time alignment, replicate filtering, gap filling, and TIC normalization. The resulting processed data was submitted to the Global Natural Product Social Molecular Network platform (GNPS)^34^ to generate a feature-based molecular network (FBMN). The molecular network was generated using the following parameters: fragment ions were removed within a ±17 Da window of the precursor *m/z*, precursor ion and fragment ion mass tolerance were set to 0.02 Da and edges were filtered to have a score above 0.7 and at least 4 matched peaks. Edges were kept if both nodes were present in each other’s top 10 most similar nodes, and molecular families’ maximum size was set to 100. Experimental fragmentation spectra were searched against GNPS’s spectral libraries and filtered in the same way (cosine score above 0.7 and a minimum of 4 matched peaks). CMMC enrichment downstream analysis was performed using default parameters. FBMN and CMMC enrichment were visualized using Cytoscape^58^ and the CMMC analysis dashboard.

## References

1. Quinn, R. A. et al. Global chemical effects of the microbiome include new bile-acid conjugations. Nature 579, 123–129 (2020).

2. Tremaroli, V. & Bäckhed, F. Functional interactions between the gut microbiota and host metabolism. Nature 489, 242–249 (2012).

3. Oliphant, K. & Allen-Vercoe, E. Macronutrient metabolism by the human gut microbiome: major fermentation by-products and their impact on host health. Microbiome 7, 91 (2019).

4. Hadadi, N., Berweiler, V., Wang, H. & Trajkovski, M. Intestinal microbiota as a route for micronutrient bioavailability. Curr. Opin. Endocr. Metab. Res. 20, 100285 (2021).

5. Wilson, I. D. & Nicholson, J. K. Gut microbiome interactions with drug metabolism, efficacy, and toxicity. Transl. Res. 179, 204–222 (2017).

6. Chiu, K., Warner, G., Nowak, R. A., Flaws, J. A. & Mei, W. The impact of environmental chemicals on the gut microbiome. Toxicol. Sci. 176, 253–284 (2020).

7. Lindell, A. E., Zimmermann-Kogadeeva, M. & Patil, K. R. Multimodal interactions of drugs, natural compounds and pollutants with the gut microbiota. Nat. Rev. Microbiol. 20, 431–443 (2022).

8. Cryan, J. F. et al. The Microbiota-gut-brain axis. Physiol. Rev. 99, 1877–2013 (2019).

9. Corbin, K. D. et al. Host-diet-gut microbiome interactions influence human energy balance: a randomized clinical trial. Nat. Commun. 14, 3161 (2023).

10. Han, S. et al. A metabolomics pipeline for the mechanistic interrogation of the gut microbiome. Nature 595, 415–420 (2021).

11. Shiroma, H. et al. Enteropathway: the metabolic pathway database for the human gut microbiota. Brief. Bioinform. 25, (2024).

12. Wishart, D. S. et al. MiMeDB: The Human Microbial Metabolome Database. Nucleic Acids Res. 51, D611–D620 (2023).

13. Kruger, R. et al. MiMeDB 2.0: The human Microbial Metabolome Database for 2026. Nucleic Acids Res. (2025) doi:10.1093/nar/gkaf1272.

14. Wortmann, E., Adam, G. & Limonciel, A. Biocrates. MxP® Quant 1000 in microbiome research. Preprint at https://biocrates.com/wp-content/uploads/2025/05/Application-note-Quant-1000-in-microbiome-research.pdf (2025).

15. Zuffa, S. et al. microbeMASST: a taxonomically informed mass spectrometry search tool for microbial metabolomics data. Nat Microbiol 9, 336–345 (2024).

16. Wang, M. et al. Sharing and community curation of mass spectrometry data with Global Natural Products Social Molecular Networking. Nat. Biotechnol. 34, 828–837 (2016).

17. Caraballo-Rodríguez, A. M. et al. The undiscovered natural product potential of Actinomycetes. J. Antibiot. (Tokyo) (2025) doi:10.1038/s41429-025-00876-x.

18. Poynton, E. F. et al. The Natural Products Atlas 3.0: extending the database of microbially derived natural products. Nucleic Acids Res. 53, D691–D699 (2025).

19. Zhao, H. N. et al. A resource to empirically establish drug exposure records directly from untargeted metabolomics data. Nat. Commun. 16, 10600 (2025).

20. Patan, A. et al. Charting the undiscovered metabolome with synthetic multiplexing. bioRxiv (2025) doi:10.1101/2025.11.18.689170.

21. Mannochio-Russo, H. et al. The microbiome diversifies long-to short-chain fatty acid-derived N-acyl lipids. Cell (2025) doi:10.1016/j.cell.2025.05.015.

22. Mannochio-Russo, H. et al. Bridging complexity and accessibility in metabolomics with MetaboApps. ChemRxiv (2025) doi:10.26434/chemrxiv-2025-3nq29.

23. Wu, M. et al. Gut complement induced by the microbiota combats pathogens and spares commensals. Cell 187, 897–913.e18 (2024).

24. Won, T. H. et al. Host metabolism balances microbial regulation of bile acid signalling. Nature 638, 216–224 (2025).

25. Nothias, L.-F. et al. Feature-based molecular networking in the GNPS analysis environment. Nat. Methods 17, 905–908 (2020).

26. Watrous, J. et al. Mass spectral molecular networking of living microbial colonies. Proc. Natl. Acad. Sci. U. S. A. 109, E1743–52 (2012).

27. McDonald, D. et al. American gut: An open platform for citizen science microbiome research. mSystems 3, (2018).

28. Mohanty, I. et al. The changing metabolic landscape of bile acids -keys to metabolism and immune regulation. Nat. Rev. Gastroenterol. Hepatol. 21, 493–516 (2024).

29. Jia, M. et al. Gut microbiota dysbiosis promotes cognitive impairment via bile acid metabolism in major depressive disorder. Transl. Psychiatry 14, 503 (2024).

30. Fogelson, K. A., Dorrestein, P. C., Zarrinpar, A. & Knight, R. The gut microbial bile acid modulation and its relevance to digestive health and diseases. Gastroenterology 164, 1069–1085 (2023).

31. Mann, A. et al. Palmitoyl Serine: An Endogenous Neuroprotective Endocannabinoid-Like Entity After Traumatic Brain Injury. J. Neuroimmune Pharmacol. 10, 356–363 (2015).

32. Long, J. Z. et al. The secreted enzyme PM20D1 regulates lipidated amino acid uncouplers of mitochondria. Cell 166, 424–435 (2016).

33. Wan, K. X., Vidavsky, I. & Gross, M. L. Comparing similar spectra: from similarity index to spectral contrast angle. J. Am. Soc. Mass Spectrom. 13, 85–88 (2002).

34. Wilkinson, M. D. et al. The FAIR Guiding Principles for scientific data management and stewardship. Sci Data 3, 160018 (2016).

35. van Santen, J. A. et al. The Natural Products Atlas 2.0: a database of microbially-derived natural products. Nucleic Acids Res. 50, D1317–D1323 (2022)

36. van Santen, J. A. et al. The Natural Products Atlas: An open access knowledge base for microbial natural products discovery. ACS Cent. Sci. 5, 1824–1833 (2019).

37. Di Tommaso, P. et al. Nextflow enables reproducible computational workflows. Nat. Biotechnol. 35, 316–319 (2017).

38. Bittremieux, W. et al. Universal MS/MS Visualization and Retrieval with the Metabolomics Spectrum Resolver Web Service. bioRxiv 2020.05.09.086066 (2020) doi:10.1101/2020.05.09.086066.

39. Kim, H. W. et al. NPClassifier: A deep neural network-based structural classification tool for natural products. J. Nat. Prod. 84, 2795–2807 (2021).

40. Lex, A., Gehlenborg, N., Strobelt, H., Vuillemot, R. & Pfister, H. UpSet: Visualization of intersecting sets. IEEE Trans. Vis. Comput. Graph. 20, 1983–1992 (2014).

## References

1. Wilson, I. D. & Nicholson, J. K. Gut microbiome interactions with drug metabolism, efficacy, and toxicity. Transl. Res. 179, 204–222 (2017).

2. Zhao, H. N. et al. Empirically establishing drug exposure records directly from untargeted metabolomics data. bioRxiv (2024) doi:10.1101/2024.10.07.617109.

3. Carmody, R. N. & Turnbaugh, P. J. Host-microbial interactions in the metabolism of therapeutic and diet-derived xenobiotics. J. Clin. Invest. 124, 4173–4181 (2014).

4. Brennan, C. et al. Clearing the plate: a strategic approach to mitigate well-to-well contamination in large-scale microbiome studies. mSystems 9, e0098524 (2024).

5. Kennedy, E. A., King, K. Y. & Baldridge, M. T. Mouse Microbiota models: Comparing germ-free mice and antibiotics treatment as tools for modifying gut bacteria. Front. Physiol. 9, 1534 (2018).

6. Zimmermann, M., Zimmermann-Kogadeeva, M., Wegmann, R. & Goodman, A. L. Mapping human microbiome drug metabolism by gut bacteria and their genes. Nature 570, 462–467 (2019).

7. Rowland, I. et al. Gut microbiota functions: metabolism of nutrients and other food components. Eur. J. Nutr. 57, 1–24 (2018).

8. Quinn, R. A. et al. Global chemical effects of the microbiome include new bile-acid conjugations. Nature 579, 123–129 (2020).

9. Cheng, J., Venkatesh, S., Ke, K., Barratt, M. J. & Gordon, J. I. A human gut Faecalibacterium prausnitzii fatty acid amide hydrolase. Science 386, eado6828 (2024).

10. Tamanna, I. et al. Microbial lipid shifts in a multi-stage simulated gut. bioRxivorg (2025) doi:10.1101/2025.09.25.678496.

11. Mannochio-Russo, H. et al. The microbiome diversifies long-to short-chain fatty acid-derived N-acyl lipids. Cell (2025) doi:10.1016/j.cell.2025.05.015.

12. Cohen, L. J. et al. Commensal bacteria make GPCR ligands that mimic human signalling molecules. Nature 549, 48–53 (2017).

13. Sánchez-Baracaldo, P., Bianchini, G., Wilson, J. D. & Knoll, A. H. Cyanobacteria and biogeochemical cycles through Earth history. Trends Microbiol. 30, 143–157 (2022).

14. Chen, M.-Y. et al. Comparative genomics reveals insights into cyanobacterial evolution and habitat adaptation. ISME J. 15, 211–227 (2021).

15. Jones, M. R. et al. CyanoMetDB, a comprehensive public database of secondary metabolites from cyanobacteria. Water Res. 196, 117017 (2021).

16. Weiss, M. B. et al. Chemical diversity of cyanobacterial natural products. Nat. Prod. Rep. 42, 6–49 (2025).

17. Janssen, E. M.-L. et al. S75 | CyanoMetDB | Comprehensive database of secondary metabolites from cyanobacteria. Zenodo 10.5281/ZENODO.13854577 (2024).

18. Huang, I.-S. & Zimba, P. V. Cyanobacterial bioactive metabolites-A review of their chemistry and biology. Harmful Algae 83, 42–94 (2019).

19. Soares, R. et al. Machine learning-driven discovery and database of Cyanobacteria bioactive compounds: A resource for therapeutics and bioremediation. J. Chem. Inf. Model. 64, 9576–9593 (2024).

20. Chorus, I. & Welker, M. Toxic Cyanobacteria in Water. (CRC Press, London, 2021).

21. Wood, R. Acute animal and human poisonings from cyanotoxin exposure - A review of the literature. Environ. Int. 91, 276–282 (2016).

22. Ettoumi, A. et al. Bioaccumulation of cyanobacterial toxins in aquatic organisms and its consequences for public health. in Zooplankton and Phytoplankton: Types, Characteristics, and Ecology (Nova Science Publishers, 2011).

23. Karjalainen, M. et al. Nodularin accumulation during cyanobacterial blooms and experimental depuration in zooplankton. Mar. Biol. 148, 683–691 (2006).

24. Huisman, J. et al. Cyanobacterial blooms. Nat. Rev. Microbiol. 16, 471–483 (2018).

25. Brooks, B. W. et al. Are harmful algal blooms becoming the greatest inland water quality threat to public health and aquatic ecosystems? Environ. Toxicol. Chem. 35, 6–13 (2016).

26. Paerl, H. W. & Paul, V. J. Climate change: links to global expansion of harmful cyanobacteria. Water Res. 46, 1349–1363 (2012).

27. Visser, P. M. et al. How rising CO2 and global warming may stimulate harmful cyanobacterial blooms. Harmful Algae 54, 145–159 (2016).

28. Merder, J. et al. Geographic redistribution of microcystin hotspots in response to climate warming. Nat Water 1, 844–854 (2023).

29. He, Y. et al. Secondary metabolites from cyanobacteria: source, chemistry, bioactivities, biosynthesis and total synthesis. Phytochem. Rev. 24, 483–525 (2025).

30. Ibelings, B. W., Backer, L. C., Kardinaal, W. E. A. & Chorus, I. Current approaches to cyanotoxin risk assessment and risk management around the globe. Harmful Algae 49, 63– 74 (2015).

31. Kaloudis, T. et al. Determination of microcystins and nodularin (cyanobacterial toxins) in water by LC-MS/MS. Monitoring of Lake Marathonas, a water reservoir of Athens, Greece. J. Hazard. Mater. 263 Pt 1, 105–115 (2013).

32. Watrous, J. et al. Mass spectral molecular networking of living microbial colonies. Proc. Natl. Acad. Sci. U. S. A. 109, E1743–52 (2012).

33. Wang, M. et al. Sharing and community curation of mass spectrometry data with Global Natural Products Social Molecular Networking. Nat. Biotechnol. 34, 828–837 (2016).

34. Nothias, L.-F. et al. Feature-based molecular networking in the GNPS analysis environment. Nat. Methods 17, 905–908 (2020).

35. Konkel, R. et al. Structural diversity and biological activity of cyanopeptolins produced by Nostoc edaphicum CCNP1411. Mar. Drugs 21, (2023).

36. Liu, J. et al. Diversity, biosynthesis and bioactivity of aeruginosins, a family of Cyanobacteria-derived nonribosomal linear tetrapeptides. Mar. Drugs 21, 217 (2023).

37. Overlingė, D., Cegłowska, M., Konkel, R. & Mazur-Marzec, H. Aeruginosin 525 (AER525) from Cyanobacterium Aphanizomenon Sp. (KUCC C2): A new Serine proteases inhibitor. Mar. Drugs 22, (2024).

38. McCall, L.-I.Zhang, W.-W. & Matlashewski, G. Determinants for the development of visceral leishmaniasis disease. PLoS Pathog 9, e1003053 (2013).

39. Lewis, M. D. et al. Fatal progression of experimental visceral leishmaniasis is associated with intestinal parasitism and secondary infection by commensal bacteria, and is delayed by antibiotic prophylaxis. PLoS Pathog 16, e1008456 (2020).

40. Lesani, M., Gosmanov, C., Paun, A., Lewis, M. D. & McCall, L.-I. Impact of Visceral Leishmaniasis on Local Organ Metabolism in Hamsters. Metabolites 12, (2022).

41. Zhang, B., Kuipers, F., de Boer, J. F. & Kuivenhoven, J. A. Modulation of Bile Acid Metabolism to Improve Plasma Lipid and Lipoprotein Profiles. J Clin Med 11, (2021).

42. Ewald, S., Nasuhidehnavi, A., Feng, T.-Y., Lesani, M. & McCall, L.-I. The intersection of host metabolism and immune responses to infection with kinetoplastid and apicomplexan parasites. Microbiol Mol Biol Rev 88, e0016422 (2024).

43. Rohwer, F., Seguritan, V., Azam, F. & Knowlton, N. Diversity and distribution of coral-associated bacteria. Mar. Ecol. Prog. Ser. 243, 1–10 (2002).

44. Matthews, J. L. et al. Symbiodiniaceae-bacteria interactions: rethinking metabolite exchange in reef-building corals as multi-partner metabolic networks. Environ. Microbiol. 22, 1675–1687 (2020).

45. Ritchie, K. B. Bacterial symbionts of corals and Symbiodinium. in Beneficial Microorganisms in Multicellular Life Forms 139–150 (Springer Berlin Heidelberg, Berlin, Heidelberg, 2012).

46. Thompson, J. R., Rivera, H. E., Closek, C. J. & Medina, M. Microbes in the coral holobiont: partners through evolution, development, and ecological interactions. Front. Cell. Infect. Microbiol. 4, 176 (2014).

47. Puntin, G. et al. Harnessing the power of model organisms to unravel microbial functions in the coral holobiont. Microbiol. Mol. Biol. Rev. 86, e0005322 (2022).

48. Sweet, M. et al. Insights into the cultured bacterial fraction of corals. mSystems 6, e0124920 (2021).

49. Bove, C. B., Ingersoll, M. V. & Davies, S. W. Help me, symbionts, you’re my only hope: Approaches to accelerate our understanding of coral holobiont interactions. Integr. Comp. Biol. 62, 1756–1769 (2022).

50. Maruyama, T. et al. Multi-omics analysis reveals cross-organism interactions in coral holobiont. bioRxiv (2021) doi:10.1101/2021.10.25.465660.

51. Yao, S. et al. Microalgae-bacteria symbiosis in microalgal growth and biofuel production: a review. J. Appl. Microbiol. 126, 359–368 (2019).

52. Kapoore, R. V., Padmaperuma, G., Maneein, S. & Vaidyanathan, S. Co-culturing microbial consortia: approaches for applications in biomanufacturing and bioprocessing. Crit. Rev. Biotechnol. 42, 46–72 (2022).

53. Song, X., Ju, Y., Chen, L. & Zhang, W. Strategies and tools to construct stable and efficient artificial coculture systems as biosynthetic platforms for biomass conversion. Biotechnol. Biofuels Bioprod. 17, 148 (2024).

54. Zhang, C. & Straight, P. D. Antibiotic discovery through microbial interactions. Curr. Opin. Microbiol. 51, 64–71 (2019).

55. Knowles, S. L., Raja, H. A., Roberts, C. D. & Oberlies, N. H. Fungal-fungal co-culture: a primer for generating chemical diversity. Nat. Prod. Rep. 39, 1557–1573 (2022).

56. Hider, R. C. & Kong, X. Chemistry and biology of siderophores. Nat. Prod. Rep. 27, 637– 657 (2010).

57. Pluskal, T., Castillo, S., Villar-Briones, A. & Oresic, M. MZmine 2: modular framework for processing, visualizing, and analyzing mass spectrometry-based molecular profile data. BMC Bioinformatics 11, 395 (2010).

58. Shannon, P. et al. Cytoscape: a software environment for integrated models of biomolecular interaction networks. Genome Res. 13, 2498–2504 (2003).

